# Leveraging selective hippocampal vulnerability among Alzheimer’s disease subtypes reveals a novel tau binding partner SERPINA5

**DOI:** 10.1101/2020.12.18.423469

**Authors:** Angela M. Crist, Kelly M. Hinkle, Xue Wang, Christina M. Moloney, Billie J. Matchett, Sydney A. Labuzan, Isabelle Frankenhauser, Nkem O. Azu, Amanda M. Liesinger, Elizabeth R. Lesser, Daniel J. Serie, Zachary S. Quicksall, Tulsi A. Patel, Troy P. Carnwath, Michael DeTure, Xiaojia Tang, Ronald C. Petersen, Ranjan Duara, Neill R. Graff-Radford, Mariet Allen, Minerva M. Carrasquillo, Hu Li, Owen A. Ross, Nilufer Ertekin-Taner, Dennis W. Dickson, Yan W. Asmann, Rickey E. Carter, Melissa E. Murray

## Abstract

Selective vulnerability is a central concept to the myriad of devastating neurodegenerative disorders. Although hippocampus and cortex are selectively vulnerable in Alzheimer’s disease (AD), the degree of involvement lies along a spectrum that we previously defined as AD subtypes revealing distinct clinical correlates. To operationalize heterogeneity of disease spectrum, we classified corticolimbic patterns of neurofibrillary tangles to capture extreme and representative phenotypes. We combined bulk RNA sequencing with digital pathology to examine hippocampal vulnerability in AD. Using a multidisciplinary approach, we uncovered disease-relevant hippocampal gene expression changes. Biological relevance was prioritized using machine learning and several levels of human validation. This resulted in five genes highly predictive of neuropathologically diagnosed AD: *SERPINA5, RYBP, SLC38A2, FEM1B*, and *PYDC1*. Deeper investigation revealed SERPINA5 to be a novel tau binding partner that may represent a “tipping point” in the dynamic maturity of neurofibrillary tangles. Our study highlights the importance of embracing heterogeneity of the human brain to yield promising gene candidates as exampled by *SERPINA5*.

## Introduction

Alzheimer’s disease (AD) is a multi-factorial neurodegenerative disorder characterized under the microscope by the abnormal accumulation of two misfolded proteins, tau and amyloid-β (Aβ)^1^. While both are hallmarks of the disease, they accumulate in very different fashions. Aβ accumulation is the result of enzymatic cleavage of the amyloid precursor protein^2,3^, whereas abnormal tau accumulation occurs following post-translational modification that disrupts its utility in microtubule stabilization^4,5^. Aβ plaque deposits accumulate in the extracellular space of isocortical convexities before descending into the limbic system, diencephalon, brainstem, and lastly cerebellum according to Thal amyloid phase^6^. In contrast, abnormally folded tau accumulates intracellularly as neurofibrillary tangles in subcortical nuclei and entorhinal cortex before ascending to limbic structures, association cortices, and lastly primary cortices according to Braak tangle stage^7^. Thus, two partners in the devastation of the AD brain with divergent topographic origins intersect in the limbic system, and more specifically the hippocampus, before continuing on their unique neuroanatomic journey.

Studies demonstrate Aβ plaque pathology as predictable and rarely deviating from the well-established Thal amyloid phase^8,9^. However, we identified three AD subtypes using an objective mathematical algorithm to assess corticolimbic patterns of neurofibrillary tangle pathology: hippocampal sparing AD, typical AD, and limbic predominant AD^10-13^. Typical AD brains were representative of the expected patterns of hippocampal and cortical involvement as outlined by Braak tangle stage. In contrast, we discovered two extremes that exist outside of the Braakian-concept of neurofibrillary tangle patterns. Hippocampal sparing AD cases demonstrate selective resilience of the hippocampus relative to severely involved association cortices, whereas limbic predominant AD cases demonstrate inundation of the hippocampus relative to mildly involved association cortices (**Fig. 1a**). Further examination of their clinical relevance revealed a constellation of clinical and demographic differences that were monotonically-directed among AD subtypes, including striking differences in age at onset, sex, and *APOE* ε4 status^10-13^. Given the neuropathologic and clinical differences observed among AD subtypes, we sought to test the hypothesis that the objective classification of disease spectrum could be leveraged to uncover transcriptomic changes that underlie selective vulnerability of the hippocampus in AD. Herein, we outline a novel translational neuropathology approach for gene prioritization that, coupled with machine learning, can be used to identify biologically relevant gene expression changes throughout a disease spectrum.

**Fig 1.**
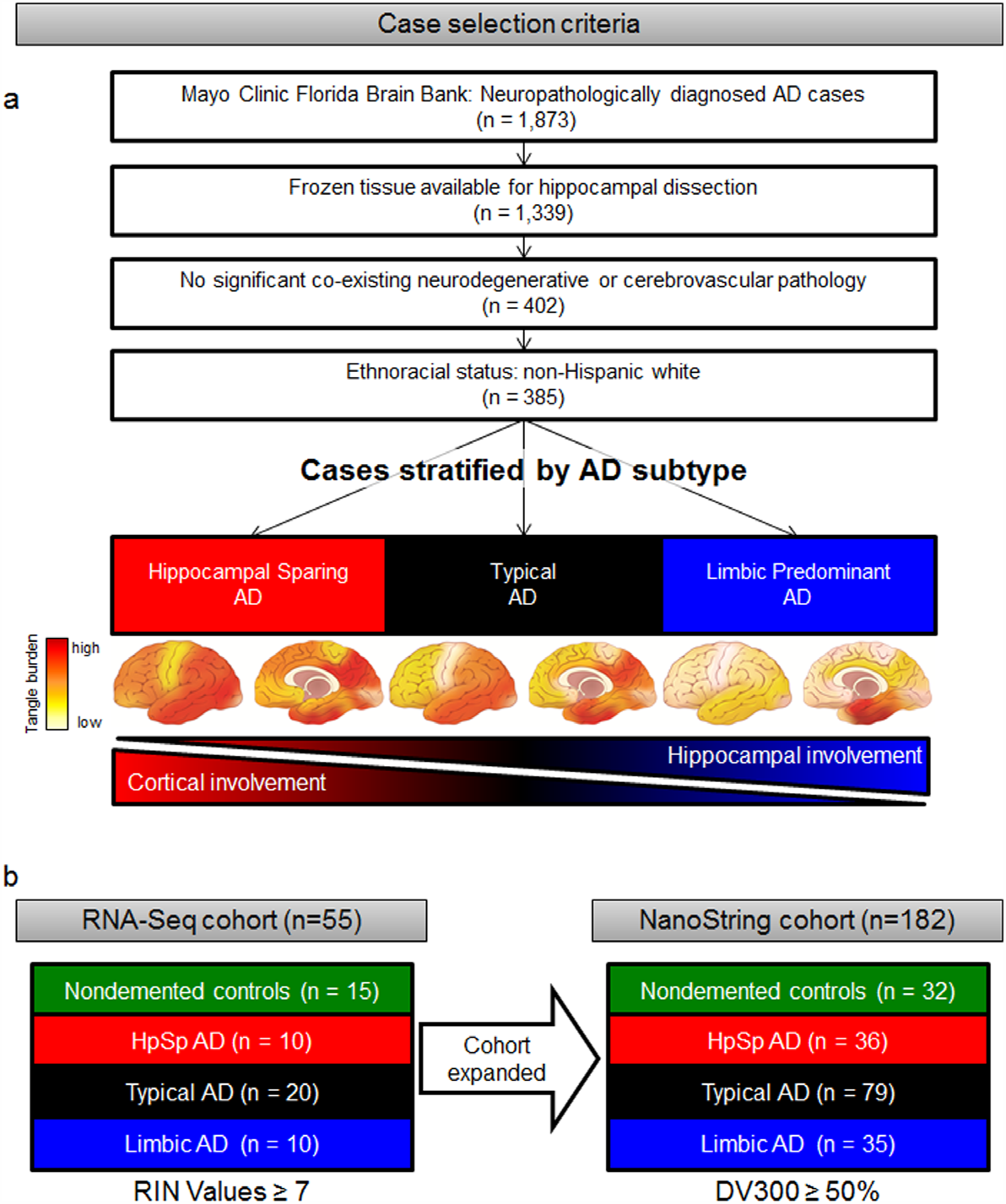
Case selection criteria and neuropathologic subtyping of AD. **a**, Flowchart describing selection criteria of AD cases to obtain frozen hippocampal tissue used to perform RNA-Seq and NanoString studies. AD was subtyped using a validated algorithm based on topographic distribution of hippocampal and cortical neurofibrillary tangles^13^. Brain cartoons illustrate corticolimbic patterns of neurofibrillary tangle burden throughout the brain in each AD subtype, where red indicates a high tangle burden and yellow/tan indicates a low tangle burden. **b**, Overview of sample size for RNA-Seq and NanoString cohorts organized by AD subtype and control classification. Note: Cohort expansion for NanoString included all RNA-Seq cohort samples. Acronyms: AD=Alzheimer’s disease, DV300=distribution value over 300

## Results

### Bulk transcriptional profiling of hippocampal vulnerability in AD

Postmortem human brain samples of neuropathologically diagnosed AD cases lacking co-existing neuropathologies and nondemented controls were derived from the Mayo Clinic brain bank (**Fig. 1a, Extended Data Fig. 1**). We obtained bulk transcriptomic data from the hippocampi of 40 AD cases and 15 controls (**Fig. 1b**) before employing a five step multi-disciplinary approach toward gene prioritization with the overall goal of selecting disease-relevant, protein-coding genes (**Fig. 2**). <u>Our first step</u> was to select genes from the literature associated with clinical expression of AD. We selected 25 genes located at loci identified in genome-wide association studies (GWAS) of late-onset AD^14-19^, 3 genes associated with early-onset AD^20,21^, and 3 additional genes identified to modify the phenotype of AD^22-24^ (**Fig. 2** [**Step 1**]). <u>Our second step</u> was to utilize our prior knowledge of disease spectrum classified by the AD subtype algorithm^13^ (**Fig. 2** [**Step 2**]). Therefore, we divided our RNA sequencing (RNA-Seq) cohort into two groups: representative phenotype (typical AD compared to control) and extreme phenotype (limbic predominant AD compared to hippocampal sparing AD) (**Fig. 2**). Given the constellation of clinicopathologic features found to differ among AD subtypes, we did not adjust RNA-Seq data by age, sex, or *APOE* ε4 status. Differential expression analysis of the representative phenotype identified 1,407 genes, among 19,720 genes that were expressed (**Fig. 2** [**Step 2a**]). Of the 1,407 differentially expressed genes in the representative phenotype, 141 genes met bioinformatic prioritization criteria and remained for further analyses of the representative phenotype (**Fig. 2** [**Step 2b**]). Differential expression analysis of the extreme phenotype identified 349 genes, among 19,747 genes that were expressed (**Fig. 2** [**Step 2a**]). Following bioinformatic prioritization, 73 genes remained for further analyses of the extreme phenotype (**Fig. 2** [**Step 2b**]). Of the 186 unique genes prioritized in **Step 2b**, only 28 (15%) overlapped between the representative phenotype and extreme phenotype.

**Fig 2.**
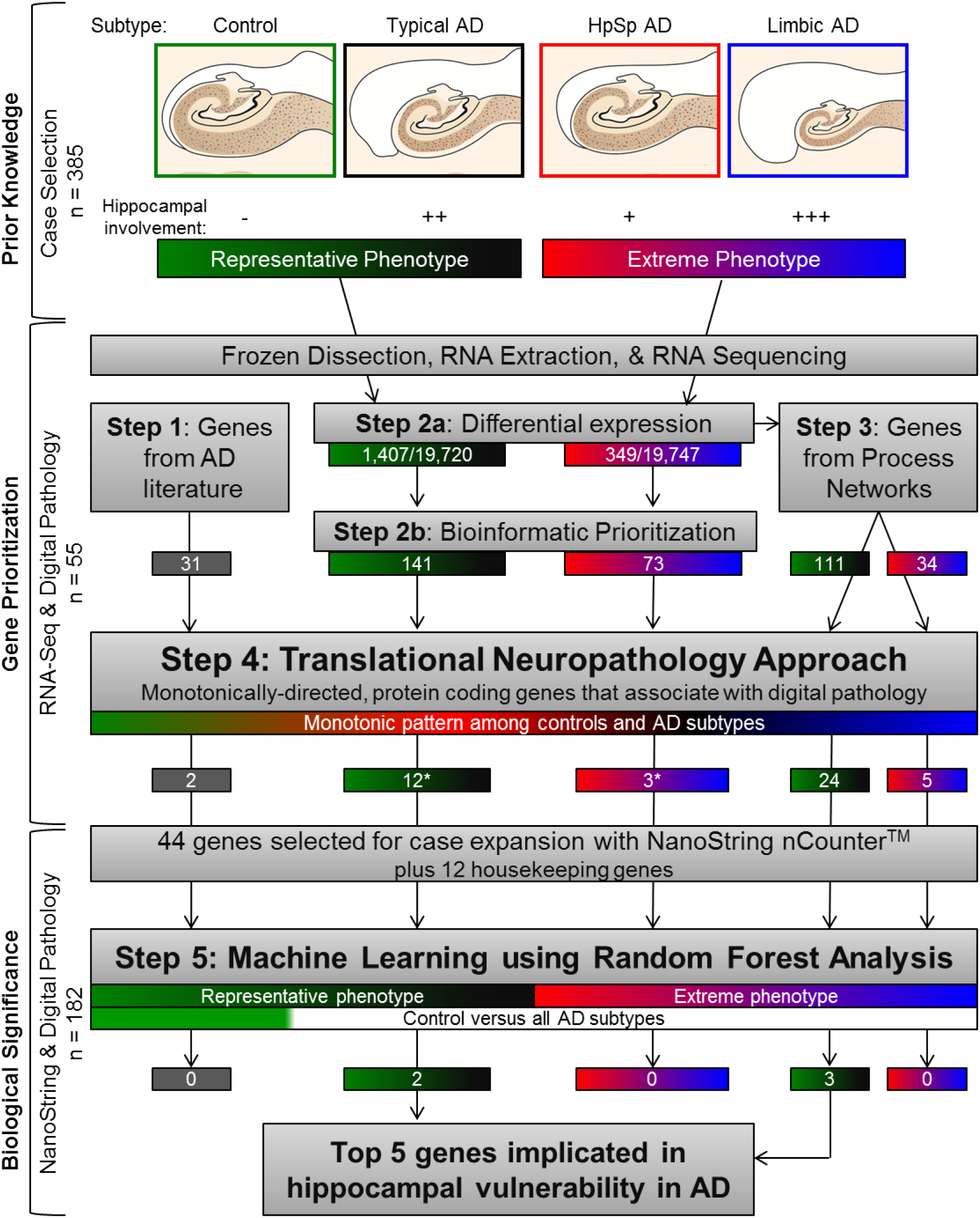
Workflow depicting novel multidisciplinary method to identify genes involved in hippocampal vulnerability in AD. Controls and typical AD cases were grouped into the representative phenotype (green-black), while hippocampal sparing AD and limbic predominant AD were grouped into the extreme phenotype (red-blue). Frozen human hippocampal tissue was first dissected, followed by RNA extraction, purification, and lastly prepped for RNA-Seq. A multi-step approach was implemented for gene prioritization, which began with genes nominated from the AD literature (**Step 1**). Next, we examined differentially expressed genes derived from the representative phenotype and extreme phenotype (**Step 2a**) before further bioinformatic prioritization (**Step 2b**). The differentially expressed genes identified in Step 2a were further prioritized based upon process network interactions (**Step 3**). We then combined the 339 unique genes from **Steps 1-3** to apply a novel translational neuropathology approach (**Step 4**) whereby protein-coding genes needed to exhibit monotonic directionality and associate with local neuropathologic markers (tau and amyloid-β) or global measures of AD pathology (Braak stage and Thal phase). Once genes were prioritized, biological significance was investigated first by expanding the RNA-Seq cohort to a total of n=182 in NanoString analyses. Machine learning was applied to NanoString data using the random forest algorithm (**Step 5**). Random forest was applied to the phenotypic groups, in addition to controls versus all AD subtypes (depicted by green vs white bar) to identify the top 5 genes predictive of hippocampal vulnerability in AD as a whole. This enabled us to utilize objective classification of disease spectrum to uncover transcriptomic changes that underlie selective vulnerability of the hippocampus in AD. **Note:** Numbers in boxes represent total gene number from each part of workflow. Steps 1-3 culminated in 339 unique genes, of which 19 overlapped in one or more of the steps. *In Step 4, one gene appeared in both groups. Acronyms: AD=Alzheimer’s disease, DE=differential expression, HpSp=Hippocampal sparing, Limbic=limbic predominant, RNA-Seq=RNA sequencing.

In both the representative phenotype and extreme phenotype, a majority of the genes were downregulated (**Fig. 3a-b**). There were 106/141 (75%) downregulated genes identified in the representative phenotype and 60/73 (82%) downregulated genes identified in the extreme phenotype. No upregulated genes were observed to overlap (**Fig. 3b**). These results suggest that within the spectrum of a disease, the pattern of upregulation or downregulation observed between disease versus controls may be recapitulated in the extremes of the disease state. In the current study, the degree of log_2_ fold change (log2FC) observed in the downregulated genes was nearly twice that observed in the upregulated genes (**Fig. 3c**).

**Fig 3.**
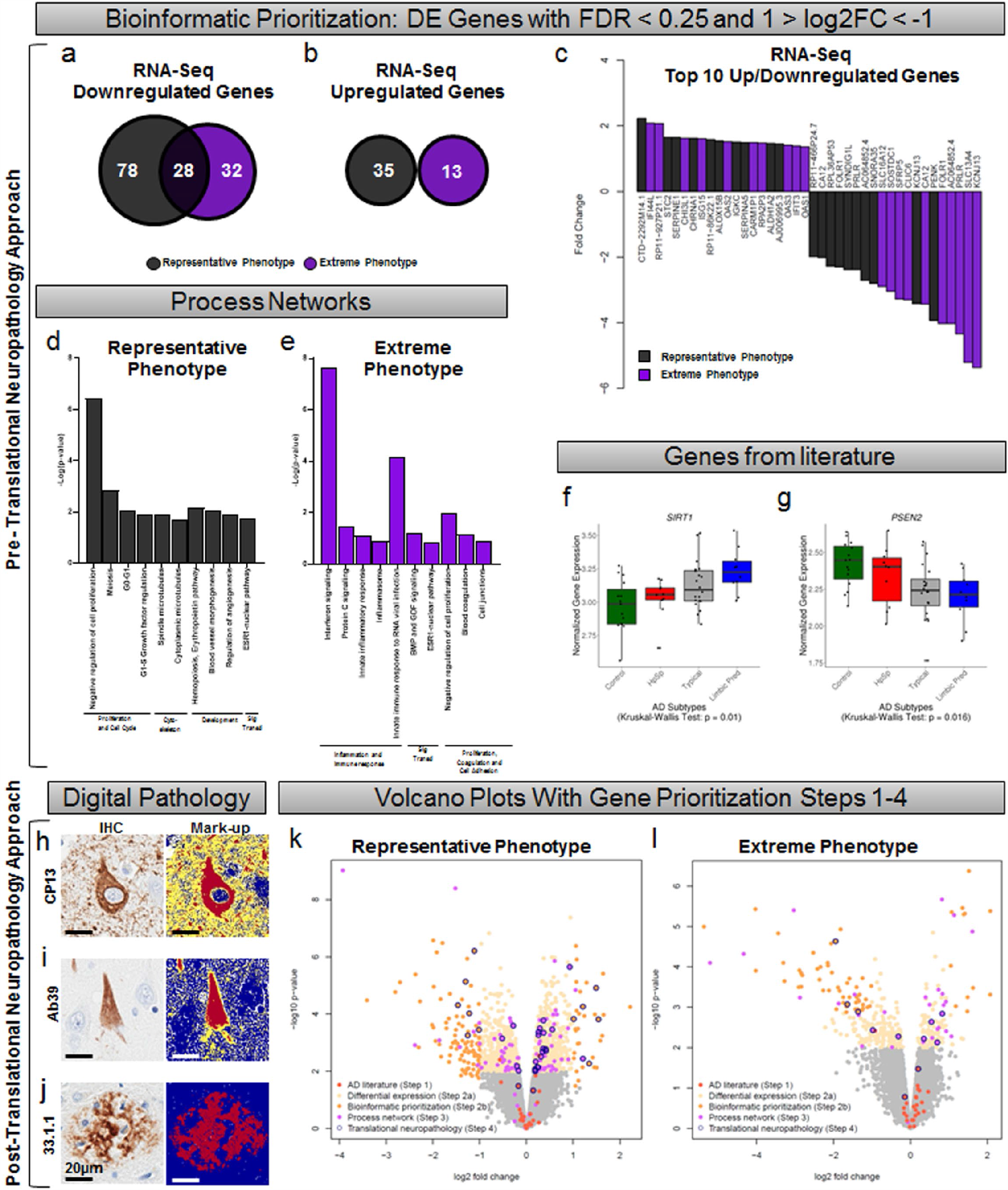
Transcriptomic analysis reveals an abundance of downregulated genes across the disease spectrum of hippocampal vulnerability in AD. **a-b**, Venn diagram depicting the number of genes that were downregulated (a) or upregulated (b) in the representative phenotype (black) or extreme phenotype (purple). There was no overlap in upregulated genes. **c**, Waterfall plot of the top ten upregulated or downregulated genes from each phenotype. **d**, Process networks in the representative phenotype (black) and extreme phenotype (purple) reveal non-overlapping functions. **f-g**, Examples of genes selected from the literature that exhibit monotonic directionality. *SIRT1* is monotonically upregulated (f) while *PSEN2* is monotonically downregulated (Statistical values can be found in **Extended Data Fig. 2**) (g). **h-j**, Digital pathology examples of immunohistochemistry and burden analysis markup using (h) CP13, (i) Ab39, and (j) 33.1.1 antibodies in CA1 subsector of the hippocampus of typical AD brains. **i-j**, Volcano plots showing fold change versus significance in the representative phenotype (i) and the extreme phenotype (j). Volcano plots

Differentially expressed genes in isolation may not be representative of their own importance to the phenotype, but they could instead reflect the indirect relationship with a gene network^25^. Thus, <u>our third step</u> was to identify differentially expressed genes from Step 2a that were enriched in pre-built process networks using a curated database created by MetaCore™ to model main cellular processes (**Fig. 2** [**Step 3**]). All genes from the top process networks, that were selected for the representative phenotype and separately for the extreme phenotype, were then included as candidate genes. The representative phenotype was highly enriched for genes associated with cell growth and proliferation, whereas the extreme phenotype was highly enriched for genes associated with inflammation and immune response (**Fig. 3d-e**). A total of 111 genes were selected from the top 10 process networks identified in the representative phenotype and 34 genes were selected from those identified in the extreme phenotype (**Fig. 2** [**Step 3**]).

### Translational neuropathology approach

Our overall goal was to identify biologically meaningful gene expression changes that relate to AD neuropathology and cognition, therefore we employed a novel translational neuropathology approach. Following steps 1-3, we had narrowed down our gene set to a total of 339 unique genes to be reviewed in <u>our fourth step</u> toward final gene prioritization (**Fig. 2** [**Step 4**]). The disease spectrum was separated back into four distinct groups to enforce monotonic directionality of gene expression changes in an expected direction of upregulation (control < hippocampal sparing AD < typical AD < limbic predominant AD) as exampled by *SIRT1* or downregulation (limbic predominant AD < typical AD < hippocampal sparing AD < control) as exampled by *PSEN2* (**Fig. 3f-g, Extended Data Fig. 2**). Of the 182 genes that showed group-wise differences in gene expression (p<0.05), 79 were monotonically directed.

Next, we investigated the relationship of the remaining 79 genes with neuropathologic burden of tau and Aβ across serial hippocampal sections for each AD case and control using digital pathology to quantify disease burden^10,11,13,26-28^. Tau accumulation inside the neuron does not occur in isolation as a static entity, but instead it accumulates along the lifespan of a neurofibrillary tangle^29,30^. To quantitatively capture the dynamic range of neurofibrillary tangle maturity, we used an early marker of maturity (CP13, **Fig. 3h**)^31^ and an advanced marker of maturity (Ab39, **Fig. 3i**)^32^. Data are presented in **Extended Data Fig. 1**. The early marker of tangle maturity in the hippocampus was not found to differ among AD subtypes (AD specific p=0.11); however the advanced marker of tangle maturity was found to differ (AD specific p=0.018). This was not Crist et al. bioRxiv 2020 surprising given the abundance of pretangle and neuritic pathology recognized by the CP13 antibody^31^; in comparison to the Ab39 antibody which recognizes advanced tangle pathology (i.e., mature tangles and extracellular tangles) that better reflects tau-induced neurodegeneration. Aβ burden was quantified with a pan-Aβ marker (33.1.1, **Fig. 3j**) to ensure the capture of all plaque types. We did not observe a difference in Aβ burden among AD subtypes (AD specific p=0.94). To complement these local measures of pathology in the hippocampus, we additionally investigated the relationship of the prioritized genes with Braak tangle stage^7^ and Thal amyloid phase^6^ as global measures of AD pathology (**Extended Data Fig. 3**).

The remaining genes to be evaluated in **Step 4** were highly enriched for protein-coding genes (72/79 [91%]), which we focused on for evaluation of neuropathologic association. Of the 72 protein-coding genes, 44 robustly associated with a measure of tau or Aβ pathology (**Extended Data Fig. 3**). We did not identify any genes in this final set that associated with the pan-Aβ marker (33.1.1), but 23/44 (52%) were associated with Thal amyloid phase. A majority of the genes were associated with tau pathology: advanced marker of tangle maturity (Ab39, 35/44 [80%]) or Braak tangle stage (13/44 [30%]). Gene selection from **Steps 1-4** are visually represented using volcano plots of log2FC versus significance for differential gene expression in the hippocampus (**Fig. 3k-l**).

### Machine learning to identify associations with AD neuropathology

To examine biological significance of the 44 prioritized genes, we expanded our gene expression studies from 55 hippocampi evaluated using RNA-Seq (**Extended Data Fig. 1**) to 190 hippocampi to be evaluated using the NanoString nCounter platform (**Extended Data Fig. 4**). The NanoString platform provided an affordable approach to multiplexing samples by allowing the user to design a custom codeset^33^. Following log_2_ transformation of normalized NanoString gene expression, we identified seven AD cases that would require a 2.5 fold adjustment with respect to housekeeping gene correction (**Extended Data Fig. 5**). An additional case was excluded due to low gene expression counts overall. These eight cases were excluded from further analyses, resulting in a final NanoString cohort of 182 AD cases and controls (**Fig. 1b**). As expected, gene expression levels derived from RNA-Seq and NanoString platforms showed a high degree of correlation (**Extended Data Fig. 6**).

**Fig 4.**
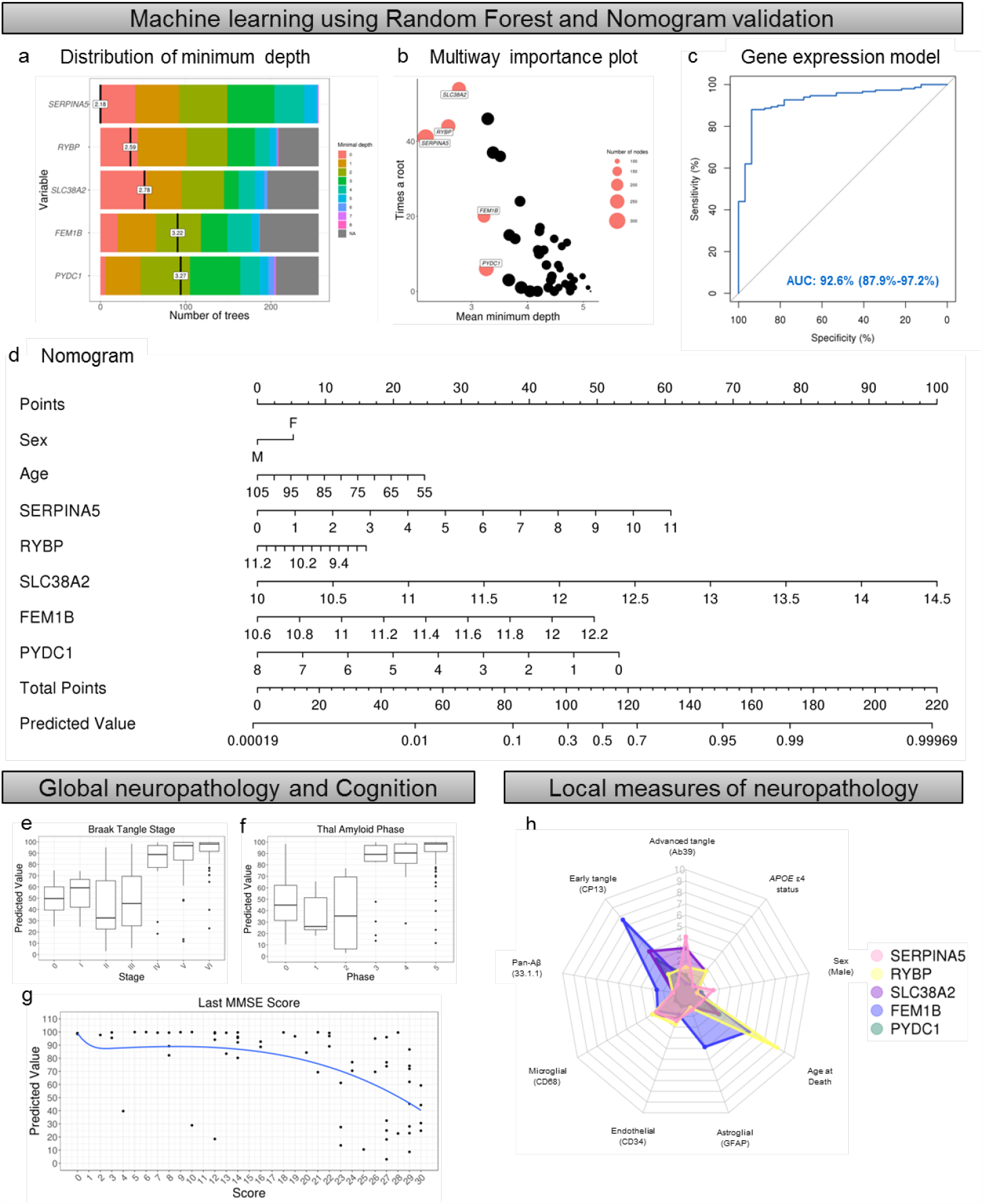
Implementation of machine learning to prioritize disease-relevant genes that associate with AD pathology and cognitive decline. **a**, Minimum depth plot generated using random forest algorithm^34^ identified the top 5 genes predictive of neuropathologically diagnosed AD based on summary of gene position within AD versus control generated trees: *SERPINA5, RYBP, SLC38A2, FEM1B, PYDC1*. **b**, Multiway importance plot depicting all genes used in random forest analysis. The top 5 are highlighted in coral, demonstrating the importance of low minimum depth (x-axis) and high root frequency (y-axis). **c**, Gene expression values were built into a logistic regression model along with age and sex to estimate the ability of the top 5 genes to discriminate between neuropathologically diagnosed AD and controls. Receiver operating characteristic reveals high discrimination with an area under the curve of 92.6%. **d**, Nomogram showing the predictive value of neuropathologically diagnosed AD for each of the top 5 genes along with age and sex. The final predictive value is determined by adding up points from each variable. See **Extended Data Fig. 10** for example of utilization. **e**, Predictive value of each case and control compared to Braak tangle stage demonstrates higher predictive values corresponding to higher stages. **f**, Predictive value of each case and control compared to Thal amyloid phase demonstrates higher predictive values corresponding to higher phases. **g**, Predictive value of each case and control with corresponding last MMSE score shows higher cognitive function corresponds with lower predictive value from nomogram. **h**, Radar plot depicting the overlay of each of the five genes regressed on demographics and digital pathology measures of tau, Aβ, and cell specific markers. Radar plot axes correspond to coefficient of partial determination for each of these regressors in the individual models. Additional statistics for radar plot can be found in **Extended Data Fig. 11**. Acronym: AD=Alzheimer’s disease, AUC=area under the curve,

**Fig 5.**
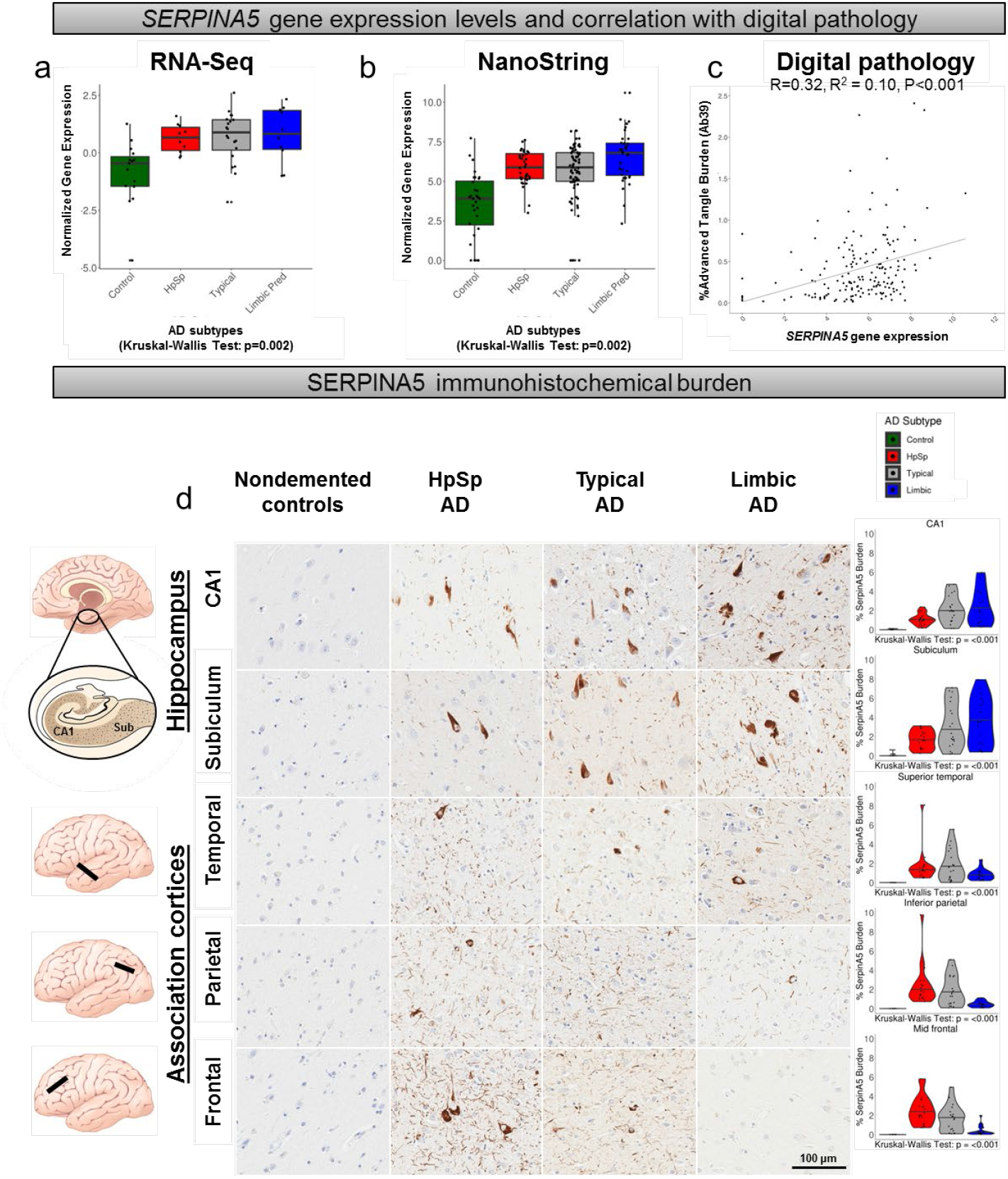
*SERPINA5* gene expression changes differ among AD subtypes and track with advanced neurofibrillary tangle pathology. **a-b**, *SERPINA5* expression increases in a monotonic fashion among AD subtypes in the RNA-Seq cohort (a), which was replicated in the larger NanoString cohort (b). **c**, *SERPINA5* gene expression from NanoString cohort associates with % burden of advanced tangle maturity marker Ab39. **d**, Immunohistochemical investigation of SERPINA5 in controls and among AD subtypes reveals a monotonic increase in hippocampal subsectors and a monotonic decrease in the association cortices. Pair-wise comparisons for graphs can be found in **Extended Data Fig. 2**. Control cases=green, Hippocampal sparing AD=red, Typical AD=grey, Limbic predominant AD=blue. Scale bar represents 100 um. Acronyms: AD=Alzheimer’s disease, Aβ=Amyloid-β, HpSp=hippocampal sparing, Limbic=limbic predominant, RNA-Seq=RNA sequencing.

**Fig 6.**
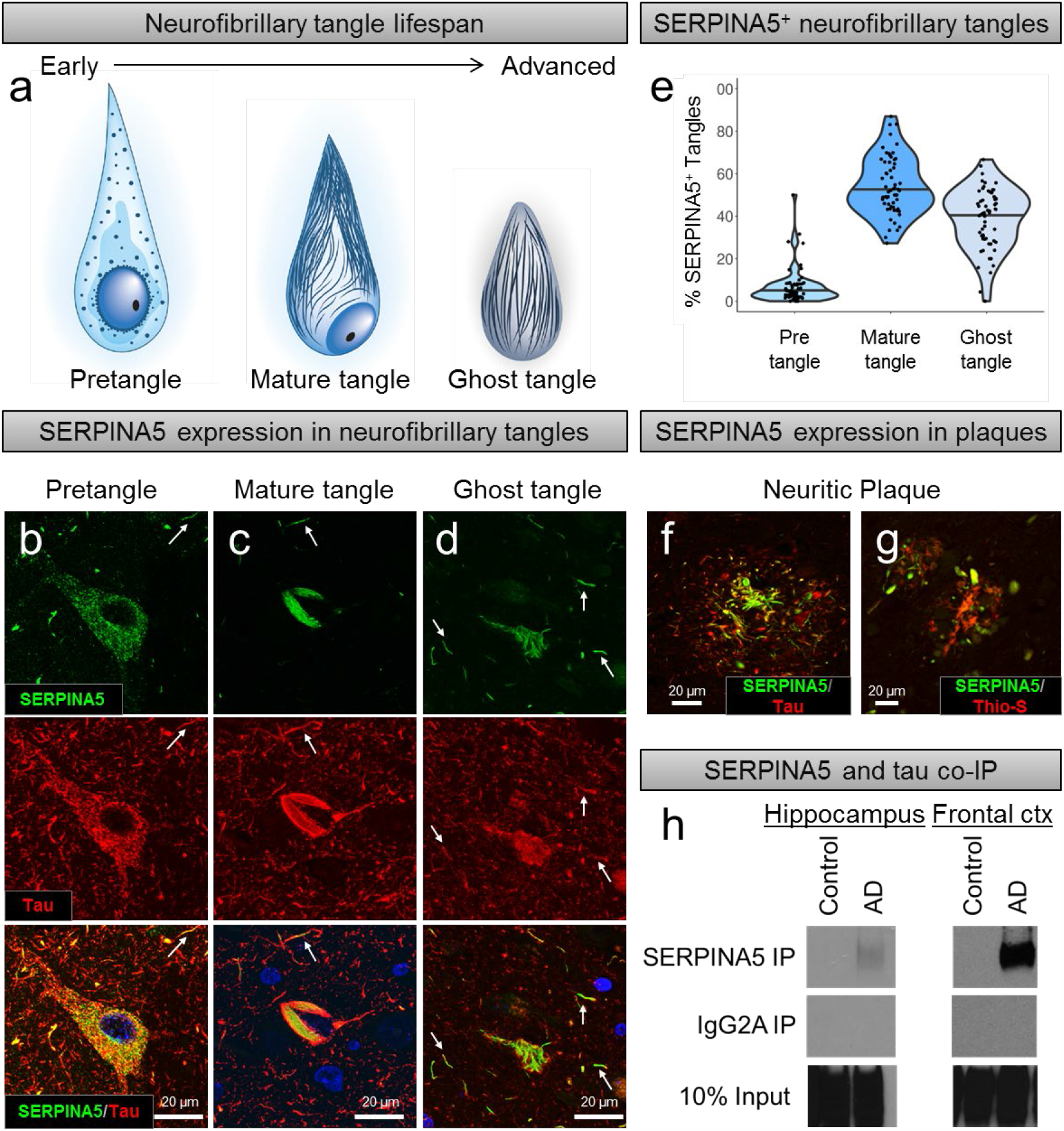
SERPINA5 is a tau binding partner expressed in neurofibrillary tangles and neuritic plaques. **a**, Cartoon depiction of the lifespan of neurofibrillary tangle maturity ranging from early to advanced forms: pretangle with punctate tau staining, mature tangle with fibrillar tau, and ghost tangle in which only the remnants of the mature tangle remain once the neuron has died. **b**, SERPINA5 accumulates alongside, as well as colocalizes with punctate tau (Tau E178, red) in pretangles. **c**, SERPINA5 accumulates in areas fairly devoid of tau and colocalizes along the fibrillar aspect of tau (Tau E178, red) in mature tangles. **d**, SERPINA5 accumulates in the extracellular space and to a lesser extent colocalizes with tau (Tau pS396, green) in ghost tangles. SERPINA5 is also observed in a subset of dystrophic neurites (arrows, b-d). **e**, Manual quantification of SERPINA5-positive neurofibrillary tangle counts in CA1 region of the posterior hippocampus shows a greater frequency of mature tangles and ghost tangles. Pretangles, mature tangles, and ghost tangles were manually quantified. **f-g**, SEPRINA5 accumulates independently and colocalizes with tau in neuritic plaques as shown by pan-tau (E1) staining and thioflavin-S. **h**, Immunoblot of pan-tau (E1) using immunoprecipitated SERPINA5 demonstrates a tau-SERPINA5 protein complex in the AD brain, but not in control brain. (Left) Tissue was sampled from frozen hippocampi of a 73 year old male control (Braak=I, Thal=0) and an 86 year old male AD case (Braak=V and Thal=5). (Right) Tissue was sampled from frozen frontal cortices of a 75 year old female control (Braak=I, Thal=0) and a 68 year old male AD case (Braak=VI and Thal=5). Note that uncropped Western blots can be found in **Extended Data Fig. 19-20**. Acronyms: AD=Alzheimer’s disease, Co-IP=Co-immunoprecipitation, Thio-S=Thioflavin-S.

Machine learning was then employed in <u>our fifth step</u> to assess the top genes predictive of hippocampal vulnerability in AD (**Fig. 2** [**Step 5**]). Our first two in silico experiments examined the top genes in the representative phenotype and in the extreme phenotype. Specifically, we employed a classification and regression method known as random forest to identify the top genes predictive for the representative phenotype and which were predictive for the extreme phenotype (see example in **Extended Data Fig. 7**). Random forest uses an ensemble of simple decision trees to classify data^34^. To facilitate prioritization and feasibility of evaluating top genes, mean minimum depth was used to rank the genes and a cutoff at the top 5 was applied (**Extended Results 1**). Out of the top 5 genes predictive when the representative phenotype and the extreme phenotype were separately analyzed, only *SERPINA5* overlapped between phenotypes (**Extended Data Fig. 8**).

The aforementioned in silico experiments provide insight into the molecular differences among AD subtypes. To maximize applicability to selective hippocampal vulnerability in AD as a whole, we combined all AD subtypes together and generated a predictive model for subsequent analysis (**Fig. 4**). Through inspection of the variable importance summaries (**Fig. 4a-b**), the following genes were considered for best discrimination of AD from control: *SERPINA5, RYBP, SLC38A2, FEM1B*, and *PYDC1*. A logistic regression model was fit to the data to estimate the ability for our top 5 genes, in concert with age and sex, to discriminate between neuropathologically diagnosed AD cases and controls. The model provided strong discrimination with an area under the curve of 92.6% (95% confidence interval [CI]: 87.9% - 97.2%; **Fig. 4c**). To represent the model visually (**Fig. 4d**), a nomogram was created^35,36^. The utility of the nomogram enables each individual’s demographic and gene expression data to be converted to a point-value (**Fig. 4d, top**), which is summarized into a total number of points that directly relates to a predicted value (**Fig. 4d, bottom**). The higher the predicted value, the more likely the data is representative of a neuropathologically diagnosed AD case. The nomogram visually provides a quantitative contribution of key factors like sex and age compared to gene expression values from the hippocampus (**Extended Data Fig. 9**). Examples of the utility of the nomogram in a nondemented control and an AD case are provided in **Extended Data Fig. 10**. The summarized predictive value was visually distinct between Braak tangle stage 0-III versus IV-VI and Thal amyloid phase 0-2 versus 3-5 when plotted (**Fig. 4 e-f**). On a clinical level, we observed a non-linear relationship between the last Mini Mental State Examination (MMSE^37^) score and nomogram predictive value, with higher MMSE scores (i.e. better cognition) corresponding to lower predicted value (**Fig. 4g**).

To examine the relationship between gene expression changes in each of the top 5 genes with local measures of neuropathology in the hippocampus, we performed a series of multiple linear regression models. Gene expression levels along with selected covariates (age, sex, and *APOE* ε4) were plotted against the following measures of neuropathologic burden: early marker of tangle maturity (CP13), advanced marker of tangle maturity (Ab39), pan-Aβ pathology (33.1.1), and cell-type specific markers to adjust for contribution of microgliosis (CD68), astrogliosis (GFAP) and endothelial burden (CD34). A neuronal marker was not included in the models to avoid regressing out genes relevant to tangle-bearing neurons. A radar plot visually displays the unique relationship between each of the prioritized genes and their degree of association with markers of neuropathologic burden (**Fig. 4h**). Each of our top 5 genes correlated with different aspects of AD pathology (results summarized in **Extended Data Fig. 11**).

### Serine protease inhibitor SERPINA5 identified as tau interactor

To validate the utility of our translational neuropathology approach to identify disease-relevant gene expression changes, we assessed the biological relevance of our top gene *SERPINA5*. To put into context of the above-mentioned steps, *SERPINA5* was first prioritized in the differential expression analysis of the representative phenotype (Step 2a: p=0.00012; Step 2b: false discovery rate[FDR]=0.0062, log2(foldchange[FC])=1.5). The translational neuropathology approach (Step 4) further identified *SERPINA5* gene expression levels to be monotonically directed (p=0.00422, **Fig. 5a**) and associate robustly with neuropathologic measures (robust association: Braak tangle stage p<0.0001 and FDR<0.10; Ab39 burden p<0.002 and FDR<0.22, and Thal amyloid phase p<0.001 and FDR<0.14, **Extended Data Fig. 3**). Random Forest analyses (Step 5, **Fig. 4a-b, Extended Data Fig. 8**) identified *SERPINA5* as the 2^nd^ top predictor in the representative phenotype (mean minimum depth=2.32, p<0.0001), 3^rd^ top predictor in the extreme phenotype (mean minimum depth=2.97, p<0.0001), and as the top predictor in the overall discrimination of AD from control (mean minimum depth=2.18, p<0.0001). Examination of three validation datasets from AMP-AD^25,38,39^ (**Extended Results 2**) revealed *SERPINA5* to be significantly upregulated in AD compared to controls in the temporal cortex (Mayo-TCX: FDR=0.000040, log2FC=2.1), superior temporal gyrus (Mount Sinai brain bank [MSBB]-Brodmann area [BM]22: FDR=0.095, log2FC=1.0), and parahippocampal gyrus (MSBB-BM36: FDR=0.0016, log2FC=2.0).

SERPINA5, also commonly known as protein C inhibitor, is a glycoprotein that inhibits serine proteases. SERPINA5 interacts with activated protein C, thrombin, and factor Xa to play a major role in hemostasis/blood coagulation and thrombosis^40-42^. Our studies revealed that in addition to its conventional role in blood coagulation, it may also play a functional role in AD pathogenesis. In both our RNA-Seq and NanoString datasets of hippocampal gene expression, we saw an increase in *SERPINA5* levels among all AD subtypes (**Fig. 5a-b**). We did not observe differences in *SERPINA5* gene expression levels when males and females were stratified within controls (p=0.92), hippocampal sparing AD (p=0.61), typical AD (p=0.83), or limbic predominant AD (p=0.87) (**Extended Data Fig. 12**). Linear regression analysis revealed *SERPINA5* expression associated with the advanced marker of tangle maturity (p=0.010) and approached significance with microgliosis (p=0.075) (**Fig. 4h, Fig. 5c, Extended Data Fig. 11**). To assess SERPINA5 localization within the brain, we utilized immunofluorescence microscopy to examine colocalization. We examined SERPINA5 immunostaining patterns across several brain cell types including astrocytes (GFAP), neurons (MAP2), microglia (resting [IBA1] and activated [CD68]), endothelial cells (CD34), and oligodendrocytes (OLIG2) (**Extended Data Fig. 13**). SERPINA5 was observed in MAP2-positive neurons in a typical AD hippocampus, whereas SERPINA5 was absent in control brain (**Extended Data Fig. 13b**). Interestingly, gene expression levels of *SERPINA5* and the neuronal marker *ENO2* inversely associate in both controls (R=-0.55, p=0.001) and AD cases (R=-0.35, p<0.001) (**Extended Data Fig. 14**).

Given SERPINA5’s association with neurofibrillary tangle pathology (**Fig. 4h**) and its predominantly neuronal expression (**Extended Data Fig. 13b**), we hypothesized that SERPINA5 would follow the corticolimbic patterns inherent to each of the AD subtypes. We employed digital pathology to quantify SERPINA5 immunohistochemical burden in hippocampal subsectors (CA1 and subiculum) and association cortices (temporal, parietal and frontal regions) vulnerable to AD neuropathologic change^7,13^ (**Fig. 5d**). Controls lacked or had minimal SERPINA5 immunopositive lesions (all medians <0.020% burden) and differed in all regions from AD cases (p≤0.002) (**Extended Data Fig. 15**). As predicted by hippocampal gene expression studies, we observed increasing immunohistochemical burden of SERPINA5 among AD subtypes in the hippocampus (AD-specific p=0.011). Based upon our observations of tangle patterns in the cortex of AD subtypes, we hypothesized we would observe an inverse monotonic directionality in the cortex. Thus, we extended our findings to investigate immunohistochemical burden of SERPINA5 across superior temporal, inferior parietal, and mid-frontal cortices (**Fig. 5d**). The average cortical burden of SERPINA5 demonstrated an overall decreasing monotonic directionality from hippocampal sparing AD > typical AD > limbic predominant AD (AD-specific p<0.001).

Next, we sought to investigate SERPINA5’s relationship with neurofibrillary tangle maturity, as more advanced tangle pathology closely associates with neuronal death^9,43^. Neurofibrillary tangles are not static entities^29,44^. They develop progressively from early pretangles with punctate tau staining and perinuclear accumulation to advanced mature tangles with fibrillar structure that form into ghost tangles with diffuse fibrils lacking a nucleus (**Fig. 6a**)^29^. To confirm the association between advanced tangle pathology (Ab39 burden) and *SERPINA5* gene expression from regression modeling, we performed immunofluorescence microscopy. In pretangles, SERPINA5, like tau, was punctate in nature and found localized to the cytoplasm and perinuclear space (**Fig. 6b**). A marked increase in SERPINA5 immunostaining was observed in mature tangles aligning with more fibrillar aspects of tau (**Fig. 6c**). Of note, areas relatively devoid of tau pathology in mature tangles appeared to be occupied by SERPINA5 accumulation. SERPINA5 immunopositive ghost tangles ranged from robustly to lightly stained in their burnt-out form as remnants of tau fibrils (**Fig. 6d**). SERPINA5 staining was predominantly found in the neuronal soma with only sparse labeling of extrasomal neuritic pathology. SERPINA5 immunopositive neurites could be found proximal to an affected neuron, which represented a subset of tau-positive neurites (**Fig. 6b-d**). Based upon our findings from immunofluorescence microscopy, we hypothesized that SERPINA5-immunopositive tangles would be in greater abundance in mature tangles and ghost tangles. The number and type of tangle were manually counted in the CA1 subsector of the hippocampus in 60 independent AD cases (**Fig. 6e**) on digitized brightfield images of SERPINA5 immunostaining. The proportion of SERPINA5-positive tangles dramatically increased in frequency from pretangles (4%) to mature tangles (52%) and ghost tangles (42%) (p<0.001). In addition to immunopositive tangles, SERPINA5 pathology was found interspersed between and colocalized with tau-positive and thioflavin-S-positive neurites in neuritic plaques (**Fig. 6f-g**).

Immunofluorescent colocalization of SERPINA5-positive/tau-positive neurofibrillary tangle pathology suggested that SERPINA5 may interact with tau. To investigate this further, we performed a co-immunoprecipitation (co-IP) experiment to determine if we detected a SERPINA5-tau protein-protein interaction. Bulk hippocampal tissue was taken from an AD case and control and used to immunoprecipitate SERPINA5, which was followed by immunoblotting using the pan-tau marker, E1^45^. We observed a strong tau signal in the total homogenate (input) of both AD cases and controls, but only detected a tau band in the SERPINA5 co-IP of the AD case (**Fig. 6h**). This initial study provided supportive evidence that SERPINA5 acts as a tau protein binding partner. To provide stronger evidence to support our hypothesis of a SERPINA5-tau interaction, we expanded our studies to the frontal cortex where larger tissue volumes can be dissected for greater yields. Western blot analysis of co-immunoprecipitation experiment revealed that tau was successfully pulled down by SERPINA5 in all three AD cases, but not in controls, suggesting that SERPINA5-tau interaction is disease-specific (**Fig. 6h**).

## Discussion

In this study, we leveraged the Alzheimer’s disease spectrum using both phenotypic differences (extreme phenotype vs. representative phenotype)^10-13^ and quantitative measures of severity (digital pathology)^10,11,26-28^ to identify key proteins involved in the molecular pathogenesis underlying selective hippocampal vulnerability. To our knowledge, this study represents the largest RNA-Seq experiment performed using bulk hippocampal tissue to date^46-48^. Using a novel approach that combined next-generation transcriptome sequencing with quantitative digital pathology and machine learning, we performed several levels of disease-relevant validation in humans that included: 1) RNA-Seq cohort expansion using NanoString technology^33^, 2) machine learning^34^ to predict utility of gene expression levels working in concert to discriminate neuropathologic classification, 3) evaluation of the relationship between gene expression levels with macroscopic scales of neuropathology (Braak^7^ and Thal^6^) and cognition (MMSE^37^), 4) digital pathology to assess biological relevance of corticolimbic distribution of SERPINA5, and 5) co-IP to examine interaction between tau and SERPINA5. In all, these tools provided invaluable insight into the complexity of hippocampal vulnerability both within the disease spectrum and in AD as a whole, highlighting the role of *SERPINA5, RYBP, SLC38A2, FEM1B*, and *PYDC1*.

Upon reviewing the published literature, our top 5 proteins were found to have a multitude of diverse functions within the living system. In addition to SERPINA5’s canonical serine protease inhibition function, it exhibits broad protease activity and interacts with phospholipids^49-53^, glycosaminoglycans^54^, proteins^55-58^, and cysteine proteases^59^. These properties make SERPINA5 extremely unique among serine protease inhibitors. RYBP is a multifunctional protein that binds several transcriptional factors^60^, mediates histone H2A monoubiquitination^61^ and plays roles in development^62,63^, apoptosis^64-66^ and cancer^60^. SLC38A2 functions as a sodium-dependent amino acid transporter that mediates both the efflux of neutral α-amino acids across the blood-brain barrier and their uptake into neurons^67-69^. FEM1B is an ankyrin repeat protein that induces replication stress signaling through its interaction with checkpoint kinase 1^70^, regulates ubiquitin-protein transferase activity^71^ and promotes apoptosis^72^. PYDC1 (also known as ASC2/POP1) regulates the innate immune response by suppressing NFκB transcription factor activity and pro-caspase-1 activation^73^.

This work underscores the importance of capturing heterogeneity as a phenotype when investigating disease-relevant gene expression changes using bulk RNA-Seq. Our ability to identify the most relevant genes was enhanced by the judicious application of the random forest algorithm^34^. This application of machine learning has wide-ranging utility outside the context of AD, as exampled by its use to uncover transcriptional signatures underlying heterogeneity of smooth muscle cells using single-cell RNA-Seq^74^. To further validate the functional relevance of our findings, we performed regression modeling to assess whether the driver of disease-relevant gene expression changes was underlying neuropathology or cellular admixture. Gene expression levels from our top gene, *SERPINA5*, were upregulated in AD and monotonically directed among AD subtypes. This is in line with a previous study that identified upregulation of *SERPINA5* in the hippocampus of late onset AD brains^46^. In addition, *SERPINA5* expression was significantly upregulated in AD in all three AMP-AD validation datasets from temporal cortex and parahippocampal gyrus^25,38,39^. The current study provides novel insight demonstrating a strong association between SERPINA5 immunopositive neurons and neuropathologic severity of advanced tangle pathology.

We sought to further investigate cellular distribution of SERPINA5 based upon a previous study describing SERPINA5 immunoreactivity in microglia^75^ and our regression model suggestive of a relationship with microgliosis. Co-immunofluorescence confirmed localization of SERPINA5 to neurofibrillary tangle-bearing neurons in AD brains. In contrast to previous findings in incidental Lewy body disease and Parkinson’s disease^75^, we did not observe protein expression in non-neural cells. This difference may be due to methodology (immunohistochemical brightfield staining versus co-immunofluorescence) or regional differences (i.e. hippocampus vs. substantia nigra). Neurons can survive the brunt of early neurofibrillary tangle pathology^76,77^, which may represent an invaluable window of time to arrest the neurodegenerative aspects of neurofibrillary tangle progression. Although, SERPINA5-immunopositive structures were only observed in the context of neurofibrillary tangle pathology, gene expression studies revealed an inverse association between *SERPINA5* and the neuronal marker *ENO2* in hippocampi of both controls and AD. This may suggest that upregulation of *SERPINA5* represents a repair process in neurons. Currently, it is unclear if SERPINA5 upregulation is a consequence of failed adaptation to neuronal dysfunction, or a contributor to neurofibrillary tangle formation. Our co-IP experiments provide supportive evidence that SERPINA5 and tau interact in the hippocampus and cortex of AD brains. This was complemented by our microscopy experiments that suggest SERPINA5 could be a tipping point between pretangles and mature tangles. Future studies will be directed at investigating the role of SERPINA5 in neurons with a focus on identifying whether SERPINA5 acts in a casual or non-causal fashion. This information could aid in identifying a mechanism to prevent or slow neuronal death.

Several limitations in the context of this research should be discussed. At the discovery stage, age and sex were not used to adjust as the AD subtypes exhibit a constellation of demographic and clinicopathologic features. We hypothesize that adjustment of these biological variables may limit identification of important gene expression changes that track with selective hippocampal vulnerability observed in the AD subtypes. As adjustment for age and sex is commonplace in bioinformatic prioritization, future studies lacking distinct clinicopathologic phenotypes should consider adjustment as appropriate to their study. The applied assumption of monotonic directionality was a reflection of the biology of the AD subtypes observed when examining clinicopathologic and demographic differences. It is important to note that the utility of this assumption may have excluded genes relevant to hippocampal vulnerability in AD that did not track with AD subtypes. The monotonic directionality of hippocampal involvement facilitated evaluation of gene expression changes from selective resilience in hippocampal sparing AD to selective vulnerability in limbic predominant AD. Although this range enabled deeper investigation into late-stage AD cases, more work is needed to determine the relevance of these gene expression changes as epiphenomenal versus active players in the pathogenesis of hippocampal vulnerability. Furthermore, the selection of literature-based genes does not include several contemporary genes, which may dramatically reduce the value of including known genes that can be leveraged using our novel approach. To provide a contemporary examination, we applied our translational neuropathology approach from Step 4 to a recently summarized gene set from Neuner et al.^78^ to identify genes that may associate with hippocampal vulnerability in AD. We identified three additional protein-coding genes that exhibited monotonic directionality among AD subtypes and robustly associated with neuropathologic measures: *ADAM10, ANKMY2, ATP5F1* (**Extended Results 3**). We encourage the research community to investigate these and other genes from our NanoString gene list in future studies. Future studies will strive to validate these genes and others prioritized for NanoString in an ethnoracially diverse series to ensure generalizability to all individuals suffering from this devastating disorder.

By embracing heterogeneity of the disease spectrum of AD^10-13,79^ and pursuing our hypotheses in the context of the human brain, we have uncovered a novel binding partner of tau – SERPINA5. Our study highlights the utility of a multidisciplinary, team science approach that should be encouraged in all disease-relevant fields. This approach may be especially useful in progressive disorders that accumulate aberrant proteins (e.g. motor neuron disease^80^, Parkinson’s disease^81^) and spectrum disorders such as diabetes that range in severity^82^. Through the use of sophisticated neuropathologic techniques, we were able to leverage our understanding of heterogeneity from the perspective of selective vulnerability in AD subtypes^10-13^. Coupled with machine learning, our study highlights the importance of multi-step prioritization efforts to yield promising gene candidates. In this era of big data generation, scientists need advanced methods to prioritize biologically relevant targets from large, heterogeneous human datasets. Here we outline a novel approach that offers several aspects of human-relevant validation and provides intriguing implications into the dynamic aspect of accumulating neurofibrillary tangle pathology that could be targeted to prevent further devastation of the AD brain.

## Supporting information

Extended Results 1

Extended Results 2

Extended Results 3

## Acknowledgements

This study was supported by the National Institute on Aging (R01 AG054449, P01 AG003949, P30 AG062677, P50 AG047266, U01 AG006786, and R01 AG034676), the Florida Department of Health, Ed and Ethel Moore Alzheimer’s Disease Research Program (6AZ01, 8AZ06, 20A22), and the Alzheimer’s Association (AARG-17-533458). We would like to thank Peter Davies for providing the CP13 antibody, Shu-Hui Yen for Ab39 antibody, Leonard Petrucelli and Casey Cook for the E1 antibody, and Pritam Das for the 33.1.1 antibody. We are grateful to Ariston L. Librero, Jo Landino, and Virginia Phillips for histological support; to Monica Castanedes-Casey for immunohistochemical support; and to our brain bank study coordinators Rachel LaPaille-Harwood and Jessica F. Tranovich. The Translational Neuropathology laboratory is especially grateful for the time and dedication of Samantha Davis. This work would not be possible without the generosity of the Gerstner Family Career Development Award, Center of Individualized Medicine at Mayo Clinic, and a kind gift from David and Frances Strawn. We thank the patients and their families for their generous brain donations to help further our knowledge of Alzheimer’s disease.

## AMP-AD Data Acknowledgements

This work was supported by National Institute on Aging [U01 AG046139, RF1 AG051504 to N.E.T]; National Institute of Neurological Disorders and Stroke [R01 NS080820 to N.E.T.]. Study data were provided by the following sources: The Mayo Clinic Alzheimer’s Disease Genetic Studies, led by Dr. Nilüfer Ertekin-Taner and Dr. Steven G. Younkin, Mayo Clinic, Jacksonville, FL using samples from the Mayo Clinic Study of Aging, the Mayo Clinic Alzheimer’s Disease Research Center, and the Mayo Clinic Brain Bank. Data collection was supported through funding by NIA grants P30 AG062677, R01 AG032990, U01 AG046139, R01 AG018023, U01 AG006576, U01 AG006786, R01 AG025711, R01 AG017216, R01 AG003949, NINDS grant R01 NS080820, CurePSP Foundation, and support from Mayo Foundation. Study data includes samples collected through the Sun Health Research Institute Brain and Body Donation Program of Sun City, Arizona. The Brain and Body Donation Program is supported by the National Institute of Neurological Disorders and Stroke (U24 NS072026 National Brain and Tissue Resource for Parkinson’s Disease and Related Disorders), the National Institute on Aging (P30 AG19610 Arizona Alzheimer’s Disease Core Center), the Arizona Department of Health Services (contract 211002, Arizona Alzheimer’s Research Center), the Arizona Biomedical Research Commission (contracts 4001, 0011, 05-901 and 1001 to the Arizona Parkinson’s Disease Consortium) and the Michael J. Fox Foundation for Parkinson’s Research. The results published here are in whole or in part based on data obtained from the AD Knowledge Portal (https://adknowledgeportal.synapse.org/). These data were generated from postmortem brain tissue collected through the Mount Sinai VA Medical Center Brain Bank and were provided by Dr. Eric Schadt from Mount Sinai School of Medicine.

## Author Contributions

M.E.M. designed and supervised the entire study. K.M.H., A.M.L., M.D.T., and M.E.M. dissected frozen brains. K.M.H. performed RNA extraction and quality control. D.W.D. performed neuropathologic evaluation and quantitative thioflavin-S counts. R.C.P., R.D., N.R.G., N.E.T. performed neurologic evaluation. A.M.L abstracted clinical information. X.W., Y.W.A., X.T., and H.L. performed all bioinformatic analysis. D.S. performed NanoString Analysis. E.R.L. and R.E.C. performed all statistical analysis. K.M.H., N.O.A., B.J.M evaluated and validated antibodies. I.F., S.A.L. and A.M.L. performed digital pathology. C.M.M. performed SERPINA5-positive tangle counts. A.M.C. designed and performed SERPINA5 experiments and data analysis. M.A., M.M.C., O.A.R. performed genotyping and provided substantial intellectual contribution. N.E.T provided funding, direction and supervision for the generation of Mayo Clinic AMP-AD data and analysis of all AMP-AD data in this manuscript. X.W, Z.S.Q, T.A.P., T.P.C., M.A. analyzed all AMP-AD data. M.E.M. and A.M.C. wrote the manuscript. All authors edited and reviewed the manuscript.

## Data Availability Statement

All requests for raw and analyzed data and related materials, excluding programming code, will be reviewed by the Mayo Clinic legal department and Mayo Clinic Ventures to verify whether the request is subject to any intellectual property or confidentiality obligations. Requests for patient-related data not included in the paper will not be considered. Any data and materials that can be shared will be released via a Material Transfer Agreement. The Accelerating Medicines Partnership (AMP-AD) data in this manuscript are available via the AD Knowledge Portal (https://adknowledgeportal.synapse.org).

## Code Availability Statement

Programming code related to the data preprocessing will be made available under the GNU General Public License version 3 upon request to Z.I.A.

## List of Abbreviations

AD: Alzheimer’s Disease
ANOVA: Analysis of variance
AUC: Area under the curve
Aβ: Amyloid-β
Bp: Basepairs
BME: beta-Mercaptoethanol
CI: Confidence interval
DAB: 3,3′-Diaminobenzidine
DAPI: 4′,6-diamidino-2-phenylindole
FC: Fold change
FDR: False discovery rate
GO: Gene ontology
HpSp AD: Hippocampal sparing subtype of Alzheimer’s Disease
H&E: Hematoxylin and eosin
Limbic AD: Limbic predominant subtype of Alzheimer’s Disease
MSBB: Mount Sinai VA Medical Center Brain Bank
MMSE: Mini mental state exam
NFT: Neurofibrillary tangle
OR: Odds ratio
PBS: Phosphate-buffered saline
PMSF: phenylmethylsulfonyl fluoride
RIN: RNA integrity number
RNA-Seq: RNA sequencing
ROC: receiver operating characteristic
TBST: tris-buffered saline with Tween-20
Thio-S: Thioflavin-S
Typical AD: Typical Alzheimer’s Disease

