## Extended data for "Leveraging selective hippocampal vulnerability among Alzheimer’s disease subtypes reveals a novel tau binding partner SERPINA5"

##### Overview of Extended Data file

**Extended Data Methods.** Description of all methods used in the manuscript with references.
**Extended Data Figure 1.** RNA-Seq cohort characteristics.
**Extended Data Figure 2.** Pair-wise comparisons of all box plots and violin plots.
**Extended Data Figure 3.** RNA-Seq gene expression level associations with neuropathologic measures.
**Extended Data Figure 4.** NanoString cohort characteristics.
**Extended Data Figure 5.** Exclusion criteria for normalized NanoString data.
**Extended Data Figure 6.** RNA-Seq and NanoString gene expression levels strongly correlate.
**Extended Data Figure 7.** Visual example of Random Forest.
**Extended Data Figure 8.** Random Forest algorithm applied to extreme and representative phenotype.
**Extended Data Figure 9.** Association between nomogram variables and likelihood of Alzheimer's Disease.
**Extended Data Figure 10.** Example of Nomogram utility.
**Extended Data Figure 11.** Linear regression modeling of gene expression and digital pathology findings
**Extended Data Figure 12.** SERPINA5 gene expression stratified by sex.
**Extended Data Figure 13.** SERPINA5 protein expression in brain cell types.
**Extended Data Figure 14.** SERPINA5 gene expression correlates with neuronal marker ENO2.
**Extended Data Figure 15.** Quantitative immunohistochemical burden of SERPINA5 measured by digital pathology.
**Extended Data Figure 16.** RNA-Seq raw read counts.
**Extended Data Figure 17.** AMP-AD RNAseq validation datasets reprocessed from Mayo Clinic and Mount Sinai brain bank.
**Extended Data Figure 18.** Antibody tables.
**Extended Data Figure 19.** Raw images of western blots from co-IP of hippocampus.
**Extended Data Figure 20.** Raw images of western blots from co-IP of frontal cortex.

### Extended Data Methods

#### Data reporting

Sample size was not predetermined using statistical methods.

#### Selection of brains from the Mayo Clinic Florida Brain Bank

A query of the Mayo Clinic Brain Bank identified 1,873 neuropathologically diagnosed AD cases with a known AD subtype[13] (**Fig. 1**). In order to perform gene expression studies, we excluded 534 brains that did not have available frozen tissue for dissection. In an effort to reduce neurobiological heterogeneity in the remaining 1,339 AD cases, we further excluded cases with co-existing pathologies (e.g. Lewy body pathology, tauopathies, tumors). Of the remaining 582 AD cases, we additionally excluded for significant cerebrovascular disease (e.g. infarcts, hippocampal ischemia). Of the remaining 402 AD cases, there were 16 AD cases that self-reported as non-Hispanic white, including 2 African-American/black decedents and 14 Hispanic/Latino decedents. One individual had an unknown ethnoracial status. These 17 AD cases were excluded as each of the AD subtypes was not represented and control brains were not available for matching. Of the remaining 385 AD cases, 68 were hippocampal sparing AD cases, 271 were typical AD cases, and 46 were limbic predominant AD cases. Given the inherent differences in age, sex, and *APOE* ε4 status among the AD subtypes[13], we matched typical cases to each subtype for age and sex prior to tissue dissection. AD cases with the highest RNA quality (RNA Integrity Number [RIN] ≥7) were selected for RNA sequencing (RNA-Seq) with nondemented controls selected to represent age distribution and sex. We identified 64 control brains with a Braak tangle stage[7] ≤III that lacked significant neurodegenerative pathology and cerebrovascular disease. Of these, 53 had available frozen tissue for dissection. Upon examination of the frozen brain tissue, 35 had available hippocampus for dissection. Following RNA extraction, three brains were additionally excluded for poor quality RNA. Of the remaining 32 controls, 15 with a RIN ≥7.0 were selected for RNA-Seq analyses.

The final RNA-Seq cohort (n=55) consisted of 10 hippocampal sparing AD, 20 typical AD, 10 limbic predominant AD and 15 controls (**Extended Data Fig. 1**). After obtaining RNA-Seq results, these studies were expanded for validation using a custom NanoString code set of genes on the following: hippocampal sparing AD (n=39), typical AD (n=81), limbic predominant AD (n=40) and nondemented controls [1](n=32). We note that eight of these cases were excluded from later analysis due to housekeeping gene correction (n=7) or low read counts (n=1). This resulted in a final NanoString cohort containing: hippocampal sparing AD (n=36), typical AD (n=79), limbic predominant AD (n=35) and nondemented controls (n=32) (**Extended Data Fig. 4**). The final NanoString cohort (n=182) included the RNA-Seq cohort to enable cross-platform evaluation (**Fig. 1**). All brains were acquired with appropriate ethical approval, and the research performed on postmortem samples was approved by the Mayo Clinic Research Executive Committee (IRB17-007585/ Bio00015595).

#### Demographics and Clinical history

Age, sex, and *APOE* ε4 status was available from the Mayo Clinic brain bank database. Clinical history abstraction was performed retrospectively from existing clinical records sent at the time of brain donation, as previously described[10,83]. The age at onset of cognitive symptoms was calculated by subtracting the date of birth from approximate date at onset and converted to years by dividing by 365.25. Similarly, disease duration was calculated by subtracting the date at onset from date of death. MMSE[37] score (0-30) and test date were abstracted to identify the score closest to death to serve as a measure of perimortem cognition. We report the last MMSE score tested within three years of death.

#### RNA Extraction, Purification and Integrity Testing

Total RNA was extracted from dissected hippocampal tissue using Trizol reagent (Thermo Fisher Scientific 15596026) according to manufacturer's instructions with all steps performed at room temperature (22°C) unless otherwise stated. 50-100 mg of frozen tissue was homogenized in 1 mL of Trizol. 200 µL of chloroform was added to the homogenate, mixed by inverting tubes vigorously for 15 sec and left to sit for 3 minutes. The tubes were centrifuged at 12,000 xg for 15 minutes at 4°C. Approximately 400 µL of aqueous layer was transferred to a clean RNase-free labeled tube containing 500 µL of 100% isopropanol, vortexed and incubated for 10 minutes to precipitate the RNA. Tubes were then centrifuged at 12,000 xg for 10 minutes at 4°C. Isopropanol supernatant was discarded and RNA pellet was washed with 1 mL of RNase-free 75% ethanol, vortexed and centrifuged at 7,500 xg for 5 minutes at 4°C. Ethanol was discarded and RNA pellet was air dried for 5-10 minutes avoiding complete drying of the pellet. 50-100 µL nuclease-free water (Ambion AM9938) was added to each tube (depending on the original wet weight of the tissue) and gently vortexed to dissolve RNA.

Total isolated RNA concentration was determined using a NanoDrop 1000 spectrophotometer (Thermo Scientific), and 30ug maximum of each RNA sample was then treated with DNase I (Qiagen 79254) digested on-column to remove contaminating genomic DNA. Specifically, 100 µL RNA sample was added to 350 µL of lysis buffer RLT/1% β-mercaptoethanol and mixed well. Then, 250 µL of 100% ethanol is added and then the entire 700 µL sample mix is pipetted onto a RNeasy Mini spin column (Qiagen #74106). The sample columns are centrifuged at ≥8000 xg for 15 sec at room temp and flow through is discarded (all spins are performed at this speed, duration and temperature). 350 µL of wash buffer RW1 is added to the spin columns and the flow through is discarded. 80 µL of the DNase/RDD working solution is directly added to the spin column membrane and incubated at room temp for 15 min, then briefly centrifuged for 5 sec. Again, 350 µL of RW1 is added to wash the columns and the flow through is discarded. 500 µL of RPE buffer is then added to the spin columns and the flow through is discarded. Again, 500 µL of RPE buffer is added to the spin columns and then they are inverted a few times to remove any remaining guanidine thiocyanate from the under the lids. The sample columns are centrifuged at ≥8000 xg for 2 minutes at room temp and flow through/collection tubes are discarded (replace with clean collection tubes and repeat drying for 1 minute then again discard flow through/collection tubes). Elute the RNA by placing the spin columns in RNAse-free 1.5 mL tubes and adding 100 µL RNase-free H2O. Incubate the samples for 2 minutes then centrifuge at 8000 xg for 1 minute at room temp. The high purity total RNA was stored at -80°C until use.

The total purified, DNase-treated RNA concentrations were determined using NanoDrop 1000 spectrophotometer (Thermo Scientific) and verified that the 260/230 ratio for samples were at or above 1.8. RNA quality was assessed by calculating RNA integrity number (RIN) and distribution values using RNA 6000 Nano Chips on the 2100 bioanalyzer (Agilent). RIN for a sample is determined by several factors, most importantly the calculation of total RNA ratio (proportion of the area under the 18S and 28S rRNA peaks to the total area under the curve) and the height of the 28S peak. RIN values range from 1–10, with 1 representing totally degraded RNA and 10 representing high quality intact RNA[84]. RIN values for our RNA-Seq cohorts were ≥7. Controls had a median RIN=7.6 (range 7.2-8.2), hippocampal sparing AD median RIN=7.3 (range 7.2-8.0), typical AD median RIN=7.4 (range 7.1-7.9) and limbic predominant AD median RIN 7.6 (range 7.2-7.8). The NanoString nCounter assay detects very low mRNA concentrations even in significantly degraded RNA samples. As a result, NanoString nCounter de-emphasizes the use of RIN and emphasizes the use of RNA distribution value (DV), which assesses the proportion of intact RNA fragments with greater than 300 nucleotides (DV300). For our NanoString nCounter studies, we followed best practices and selected cases and controls that had a DV300 >50%[85].

#### RNA Sequencing

RNA-Seq analysis was performed using 200 ng total RNA. Ribosomal RNA depletion and library preparation was performed using TruSeq Stranded Total RNA Library Prep Gold (Illumina 20020599, San Diego, CA). Quality of library was assessed using Agilent Bioanalyzer before sequencing. RNA library was sequenced at three samples per lane on the Illumina HiSeq2500 to generate 101 base pairs (bp) x 101 bp paired-end reads.

#### Bioinformatic Analysis of RNA-Seq data

MAP-RSeq[86], an integrative bioinformatics pipeline, was used to obtain gene read counts and various quality control matrices. More specifically, raw reads were aligned to human reference genome build hg19 using TopHat v2.0. Reads mapped to known genes were obtained using featureCounts program in Subread tool kit v1.4.4. The number of reads mapped to known genes is 46±17 million in these samples (**Extended Data Fig. 16**). Read count data were normalized using the R package cqn[87], taking into consideration the library size, gene length and GC content of each gene coding region. No outliers were detected according to principle component analysis and hierarchical clustering of normalized gene expression data. Source of variation analysis in Partek® Genomics Suite® software[88] identified that RIN contributed a significant proportion to gene expression variation on average. Genes with zero read counts in all samples were excluded from downstream analyses. This included 6,823 genes out of 57,773 total genes. Genes with a mean normalized expression (in scale of log2 Reads Per Kilobase of transcript, per Million mapped reads) below zero were marked as low expressing and filtered out from further analyses.

#### Selection of literature-based genes

Known genes associated with clinical expression of AD were selected in (**Fig. 2** [**Step 1**]). From large consortium efforts, we evaluated 25 genes that are located at loci identified in GWAS of late-onset AD: *ABCA7, APOE, BIN1, CD2AP, CD33, CLU, CR1, EPHA1, MS4A4E, MS4A6A, PICALM, CASS4, CELF1, DSG2, FERMT2, HLA-DRB1, HLA-DRB5, INPP5D, MEF2C, NME8, PTK2B, RIN3, SLC24A4, SORL1, ZCWPW1*[14-19]. The remaining literature-based genes included those associated with early-onset AD (*APP*, *PSEN1*, *PSEN2)*[20,21] or those implicated in modifying the phenotype of AD (*GRN*, *SIRT1*, *TOMM40)*[22-24].

A recent review by Neuner and colleagues[78], highlighted 112 genetic loci associated with AD (**Extended Results 3**). Of the 112 genes, one gene (*ACE*) had two available Ensemble IDs and 4 genes were not detected in our dataset. Of the 108 genes with Ensemble ID's available for post-hoc comparison in our RNA-Seq dataset, 3 additional genes would have passed our criteria: *ADAM10*, *ANKMY2*, and *ATP5F1*. The literature gene *PSEN2* is currently included in the study.

#### Gene differential expression and enrichment analysis for RNA-Seq data

R edgeR package[89] was used to analyze differentially expressed genes within the representative phenotype (typical AD compared to control) and within the extreme phenotype (limbic predominant AD compared to hippocampal sparing AD) (**Fig. 2** [**Step 2**]). A generalized linear model, adjusting for RIN, was applied with the underlying distribution set as a negative binomial to obtain log2FC and p-value of gene expression differences between the above-mentioned phenotypic groups. Given the biological relevance of age, sex, and *APOE* ε4 status to AD subtypes[10-13], these covariates were not used to adjust RNA-Seq data. Briefly, glmFit function from edgeR[89] was applied to fit a negative binomial generalized log-linear model using raw read counts, an offset matrix, dispersion parameters and design matrix. Next, glmLRT function was used to conduct a likelihood ratio test for differential expression. The offset matrix that incorporated effects of library size, gene length and GC contents was computed using R cqn package[87]. The gene wise dispersion parameters were estimated by edgeR functions estimateGLMTrendedDisp and estimateGLMTagwiseDisp. We used an unadjusted p-value cutoff of 0.01 to nominate top differentially expressed genes. To enable a broad set of differentially expressed genes to be selected for downstream prioritization efforts utilizing neuropathology measures, bioinformatic prioritization employed a less stringent cutoff (FDR<0.25) than current convention (i.e., FDR<0.10). To further enrich for genes found to be downregulated or upregulated by a factor of 2 or higher, log2FC >1 or <-1 cutoff was applied.

Enrichment analysis of differentially expressed genes from **Step 2a** was performed to identify enriched pathways and gene ontology (GO) terms using GeneGo MetaCore from Clarivate Analytics (**Fig. 2** [**Step 3**]). Significantly perturbed networks were identified by mapping differentially expressed genes from each phenotype onto pre-built process networks. Process networks were sorted by p-value and the top 10 statistically enriched process networks were reported.

#### Monotonic directionality and neuropathologic association of RNA-Seq data

We identified genes with a monotonically directed pattern of upregulation or downregulation as follows: control, hippocampal sparing AD, typical AD, limbic predominant AD (**Fig. 2** [**Step 4**]). The directionality was chosen based upon established evidence of hippocampal involvement, where the control is expected to have the least involvement and limbic predominant to have the most severely affected hippocampus[10-13]. An Analysis of Variance (ANOVA) with RIN as covariate was performed using Partek[88] to identify group-wise differences across all four groups. Genes were selected if the overall p-value from ANOVA was ≤0.05 and the mean expression monotonically increased or decreased along the spectrum: control, hippocampal sparing AD, typical AD, and limbic predominant AD. Gene class assignment of protein-coding was used to further prioritize monotonically-directed genes.

Using the linear model (lm) function in R, linear regression was performed to examine the relationship between gene expression and neuropathologic measures of tau and Aβ (**Fig. 2** [**Step 4**]). Specifically, gene expression was regressed on tau measures including Braak tangle stage, an early marker of tangle maturity (CP13, directed at phospho-S202), and an advanced marker of tangle maturity (Ab39, conformational epitope). Gene expression was regressed on Aβ measures including Thal amyloid phase[6] and a pan-Aβ marker (33.1.1, raised against Aβ1-16). RIN was included as a covariate for each of the above regression models and run separately for each tau and Aβ measure. Regression analyses for genes identified in **Steps 1-3** were examined across all four groups (control, hippocampal sparing AD, typical AD, limbic predominant AD) and within the representative phenotype (control, typical AD) (**Extended Data Fig. 3**). Genes found to robustly associate in both analyses were further selected for NanoString analyses.

#### NanoString nCounter™ Analysis

To investigate the biological significance of the prioritized genes in (**Fig. 2** [**Step 5**]), 150 ng of purified total RNA was used for NanoString nCounter gene expression analysis. A custom codeset of 56 genes was designed to validate RNA-Seq findings in the expanded NanoString cohort of hippocampal tissue from 190 brains. The 56 genes included the 45 genes prioritized in **Step 4** and 12 housekeeping genes. The 12 housekeeping genes were selected as they previously demonstrated stable expression across a diverse series of human tissue[90]. These included *C1orf43, CHMP2A, EMC7, GPI, PSMB2, PSMB4, RAB7A, REEP5, SNRPD3, VCP, VPS29,* as well as the traditional reference gene *GAPDH*. As one gene (*CFI*) was inadvertently left off of the custom codeset, the remaining analyses were conducted on 44 of the 45 prioritized RNA-Seq genes.

For NanoString nCounter analysis, R package OSAT[91] v1.18 was used to perform blocking randomization on 192 samples across all four groups (control, hippocampal sparing AD, typical AD and limbic predominant AD), sex (male or female), age (55-103) and RIN (4.6-8.9). Each NanoString plate had 2 X 6 wells and held 12 samples. Thus, age and RIN were stratified into 15 levels each. Samples were assigned to 15 plates in such a way that 13 plates held 12 samples each and 2 plates held 11 samples each. Samples were distributed in each plate as evenly as possible according to an objective function. The tests of independence between plates and sample variables (diagnosis, sex, age, and RIN) were not significant. Manual check also revealed that the number of subjects in any specific diagnosis group differed at most by 1 across plates, and mean values of age and RIN were similar across plates. To test technical reproducibility, we plated replicate RNA extracted from the brains of a hippocampal sparing AD and from a limbic predominant AD. Thus, 190 brains were assessed, but 192 samples were plated. Duplicate concordance was 98.4% for the hippocampal sparing AD and 98.1% for the limbic predominant AD.

NanoString gene expressions were assayed using the NanoStringNorm[92] package in R (version 3.4.2). Samples were normalized twice, first for quality control and second for background correction using:

$$c \text{ x } (\frac{m}{s})$$

where c was the count data, m was the geometric mean of the sum of the housekeeping genes across samples, and s was the sum of the housekeeping genes for a given sample. Samples with content greater than three standard deviations were removed during the first pass (quality control) and samples requiring a 2.5 fold adjustment after visual assessment were removed during the second pass (background correction). Seven cases were excluded from further analyses as they were identified to have a greater than 2.5 fold adjustment following housekeeping normalization (**Extended Data Fig. 5**). One case was excluded due to low gene expression counts overall. The final NanoString cohort consisted of 182 cases and controls.

#### Random Forest Modeling

After selection of the 44 candidate genes from **Steps 1-4**, additional prioritization of the candidate genes (**Fig. 2** [**Step 5**]) was required to facilitate further basic discovery research. Given the possibility of complex interactions among the genes we used a random forest model (**Extended Data Fig. 7**), which is a non-parametric, rules-based approach[34]. The set of classification trees was developed using the randomForest package[93], version 4.6-12, in R version 3.4.2. The default tuning parameters were accepted as the goal was to provide a preliminary investigation into how effectively the gene expression values could discriminate typical AD from controls in the representative phenotype, and discriminate limbic predominant AD from hippocampal sparing AD in the extreme phenotype. AD in the context of the final modeling was any of the three subtypes to enable a spectrum of hippocampal vulnerability to be captured. Thus, the final top 5 genes are derived from random forest analyses run to discriminate all AD cases from controls. Five top genes rather than 6 or 10 were arbitrarily chosen based on prioritization and feasibility for downstream applications, not based on statistical selection. Variable importance was assessed using the R package randomForestExplainer version 0.9[94]. This package enabled exploration of variable inclusion metrics, assessment of changes in model performance with exclusion of a variable, and interaction summaries within the data. Two variable inclusion metrics were considered of direct interest.

First, the number of times a variable was selected as a root was considered (node=0, **Extended Data Fig. 7**). This metric tabulates the number of times a variable was selected as a root across the 500 trees. A higher number suggests that among the randomly selected subsets of genes chosen to create each of the 500 trees, the gene consistently provided the highest degree of precision. The second metric, minimum depth, builds upon the number of times a gene was a root variable by considering where, in terms of depth of the tree, a gene was located. The closer they are to the root, the more important they are at the classification task. The minimum depth tabulates all the different positions each variable takes across all the trees in the forest. A smaller number indicates that the variable was chosen closer to the root if it was not the root. These two metrics can be plotted as a scatter plot (multiway importance plot) to understand how genes may interact. While many of the genes were found to be associated with AD status using the random forest approach (**Extended Results 1**), the top 5 genes with smallest minimum depth were selected to examine biological relevance.

#### Random Forest Model Discrimination and Nomogram

In the previously described random forest model, only gene expression values were used as candidate AD classification variables. To better understand how discriminatory the prioritized genes were in classifying AD from control, a series of multiple logistic regression models were considered. The base model was the gene expression value only. A second set of models provided adjustment for age and sex. Odds ratios (OR) and area under the receiver operating curve (AUC) were used to quantify the degree to which the variables discriminated AD from control. To present the model results succinctly in a visual form (**Fig. 4d**), a nomogram was developed[35,36]. This visual model allows one to assess relative importance of variables from the scale of the points associated with observed values and to quickly perform the non-linear calculation of probabilities using simple addition. To estimate the probability of an AD case having a higher predicted value from the nomogram compared to a control case, we evaluated a receiver operating characteristic (ROC) curve (**Fig. 4c**). We applied a conservative cutoff of 77.025, given the greater proportion of AD cases (n=150) compared to controls (n=32). This cutoff value provides the 'line' at which the model performs the best in terms of Sensitivity, Specificity, and Accuracy when classifying AD vs Control. At 77.025, our Sensitivity is 0.88 (95% CI: 0.83-0.93), our Specificity is 0.94 (95% CI: 0.85-1), and our Accuracy is 0.89 (95% CI: 0.889-0.891).

#### AMP-AD RNA-Seq Validation Datasets

We downloaded gene read counts and metadata files of three RNA-Seq cohorts from AD knowledge portal on Synapse (www.synapse.org**, Extended Data Fig. 17a**), each consisting of postmortem brains from AD and controls (**Extended Data Fig. 17b**). These included the Mayo Clinic temporal cortex (Mayo-TCX)[25,38], and Mount Sinai VA Medical Center Brain Bank superior temporal gyrus (MSBB-BM22) and parahippocampal gyrus (MSBB-BM36)[39]. The gene read counts were generated through a consensus bioinformatics pipeline that aligned raw RNA-Seq data files, counted reads mapped to genes, and reported quality control measures. Further details can be found on the AD knowledge portal (https://adknowledgeportal.synapse.org, Synapse ID: syn17010685) and by Wan et al.[95]. Neuropathologic information provided in the available metadata files was used to assign individuals as AD, control, or other (**Extended Data Fig. 17b**). AD cases and controls were the focus of our differential expression analyses. As previously described in the "Gene differential expression and enrichment analysis for RNA-Seq data" section, we performed differential expression analysis between AD and control samples using R edgeR package[89] and cqn package[87]. The covariates included in the design matrix were diagnosis (AD or control, categorical), RIN (continuous), age at death (continuous), sex (categorical) and flowcell (categorical) for all four datasets (including current study), plus the source of samples (categorical) for Mayo-TCX dataset. The resulting sample sizes for Mayo-TCX was n=68 controls and n=80 AD, MSBB-BM22 was n=33 controls and n=70 AD, MSBB-BM36 was n=30 controls and n=56 AD, and the current study was n=15 controls and n=20 typical AD (**Extended Data Fig. 17c, Extended Results 2**).

#### Immunohistochemistry

Routine sampling of the posterior hippocampus was performed at the level of the lateral geniculate at the time of brain cutting[96,97]. Formalin-fixed tissue was paraffin-embedded for archival purposes. Serial sections of the posterior hippocampal tissue blocks were cut at 5-μm and mounted to glass slides. Slides were deparaffinized using three five minute xylene washes followed by two three minute 100% ethanol washes and one three minute 95% ethanol wash. Immunohistochemical staining of tau, Aβ, and cellular markers was performed on a Dako Autostainer (Universal Staining System, Carpinteria, CA) using 3,3′-diaminobenzidine (DAB) as chromogen. Antibodies to an early marker of tangle maturity (CP13)[31], an advanced marker of tangle maturity (Ab39)[98], pan-Aβ (33.1.1), astrocytic marker (GFAP), endothelial marker (CD34), and activated microglia (CD68) were used (**Extended Data Fig. 18a**).

#### Digital pathology

Digital pathology methods were previously described in detail[10,11,13,26]. Briefly, microscope slides stained with the antibodies outlined in **Extended Data Fig. 18a** were digitally scanned using an Aperio AT2 system (Leica Biosystems, Buffalo Grove, IL) and annotated using Aperio ImageScope (Leica Biosystems, Buffalo Grove, IL, version 12.4.2.7000). The hippocampus proper was traced to enable direct comparison with gene expression data from the frozen hippocampus dissected from the contralateral side. Slides were analyzed with custom-designed macros using the color deconvolution algorithm (CP13, CD68) or positive pixel count algorithm

(Ab39, GFAP, CD34, 33.1.1). Data obtained from the color deconvolution and positive pixel count analysis was a percent burden that represented the percent area of staining out of the area annotated.

#### SERPINA5 immunohistochemical validation

Following identification of *SERPINA5* as our top gene, biological relevance was investigated immunohistochemically in an independent series of 10 controls, 20 hippocampal sparing AD, 20 typical AD, and 20 limbic predominant AD cases (**Extended Data Fig. 15**). The posterior hippocampus, superior temporal cortex, inferior parietal cortex, and mid-frontal cortex were cut in 5 μm sections and immunostained with a SERPINA5 antibody (1:100, mouse IgG, R&D Systems, Minneapolis, MN) using the Lab Vision Autostainer 480S (Thermo Scientific, Waltham, MA) and DAB (Dako, Carpinteria, CA) as chromogen. Immunostained sections were similarly digitized with Aperio technologies. The pyramidal layer of the CA1 and subiculum within the hippocampus were traced separately for analysis. The lacunosum layer marked the superior boundary and the alveus indicated the inferior boundary for tracing. Additionally, the midway of the dentate gyrus was used to operationalize the CA1-subiculum border. The cortical regions were traced in the area with the most pathology along the strait of the gyrus. To analyze the traced regions, a color deconvolution algorithm was custom-designed based upon the tinctorial properties of immunopositive lesions and set conservatively to avoid quantifying lipofuscin pigment. Data obtained was a percent burden of SERPINA5 per total area annotated.

#### SERPINA5 tangle counts

Using Aperio's Image Scope (version 12.4.3.7001), the CA1 region of the hippocampus was outlined as stated in the digital pathology methods section. The counter tool was used to mark tangle-bearing neurons in the 20 hippocampal sparing AD, 20 typical AD, and 20 limbic predominant AD cases used above. Tangle maturity levels were identified based on the following set of rules: 1) pretangles exhibit diffuse/granular staining, stain less intensely than mature tangles, and contain a nucleus; 2) mature tangles exhibit intense immunostaining (generally throughout the entire neuron) and contain a nucleus; 3) ghost tangles exhibit less intense immunostaining than mature tangles, loosely arranged bundles of fibers, and no nucleus.

As tangles occur over a lifespan of maturity[29], we also categorized two intermediary stages: 1) "Intermediary 1" consisted of the level between pretangles and mature tangles. These tangles have intense diffuse or focal granular immunostaining, as well as some fibrillary staining pattern and contained a nucleus. 2) "Intermediary 2" is the level between mature tangles and ghost tangles and was identified by intense immunostaining with no nucleus. If tangles did not fit nicely into the above categories based on shape, immunostaining, or nuclear position, these tangles were considered "unclassified" and were not included in analysis. Furthermore, tangles were not counted if the nucleus lay outside the annotated region or if less than half of the body of the tangle lay inside the annotated region. Tangles were reviewed with neuropathologists (DWD, MEM) and edited for all samples.

For analysis, Intermediary 1 counts were combined with mature tangles counts since they did not meet the requirements for pretangles. Similarly, Intermediary 2 counts were added to ghost tangles counts since they did not meet the requirements for mature tangles. Total tangles represent the number of pretangles, mature tangles, and ghost tangles in the annotated CA1 region. Percent pretangles, mature tangles, and ghost tangles were then calculated by multiplying by 100.

#### Immunofluorescent staining

Formalin-fixed, paraffin embedded posterior hippocampus sections were deparaffinized as described in the previous section. Antibody retrieval was performed by steaming slides for 30 minutes in either deionized water or citrate buffer (Dako S1699). Tissue was blocked in Dako protein block (Dako X0909) for 1 hour at room temperature. Primary antibody was diluted in Dako diluent (Dako S3022) at appropriate concentration (see **Extended Data Fig. 18b**) and allowed to incubate at 4ºC overnight. The slides were washed three times with

PBS + 0.1% Tween-20 then incubated in appropriate secondary plus DAPI (1:500, Thermo D1306) for 4 hours at room temperature. After secondary incubation (see **Extended Data Fig. 18c**), slides were washed three times with PBS + 0.1% Tween-20 and autofluorescence was quenched using 1% Sudan Black B (Sigma 199664) for 5 minutes. Slides were coverslipped using AquaPolyMount (Polysciences 18606) and imaged using a LSM 880 confocal microscope with AiryScan (Carl Zeiss Microscopy).

#### Co-Immunoprecipitation and western blot

Experiments were performed using Dynabeads Co-IP kit (Invitrogen 1432D). Posterior hippocampus from frozen brains was dissected and weighed. Tissue was then homogenized in IP buffer (IP buffer plus 100 mM NaCl, 25 mM MgCl2, 1 mM DTT, 1:200 PMSF) and allowed to incubate for 15 minutes. After spinning down at 2600 xg for 5 minutes, samples were brought to a final concentration of 0.11 g/mL. Beads were conjugated to SERPINA5 (R&D System MAB1266) antibody. Antibody-conjugated beads were then added to the homogenate and allowed to incubate for 1 hour at 4ºC. Beads were washed and sample was eluted as per manufacturer instructions.

Samples were mixed in a 3:1 ratio with Laemmli buffer (Bio-Rad 161-0747) with 10% BME (Sigma M6250) and boiled at 90ºC for 10 minutes. Samples were loaded onto a 10% Mini-PROTEAN®TGX™ Precast Protein Gels (Biorad 4561034) as well as 10 µL of precision plus dual color standards ladder (Biorad 1610374). Gel was run at 100V for 2 hours on ice. Transfer was performed using Trans-Blot® Turbo™ RTA Mini Nitrocellulose Transfer Kit (Biorad 1704158). Membrane was blocked using 5% dry milk in TBST for 1 hour before incubating in primary antibody diluted in 5% dry milk in TBST overnight at 4C. Membrane was washed several times using TBST then incubated in secondary antibody for one hour at room temperature. Secondary antibodies were as follows: Peroxidase AffiniPure $F(ab')_2$ Fragment Donkey Anti-Rabbit IgG (H+L) (Jackson Immunoresearch 711-036-152) or Peroxidase AffiniPure $F(ab')_2$ Fragment Donkey Anti-Mouse IgG (H+L) (Jackson Immunoresearch 715-036-150). Membranes were treated with SuperSignal West Pico Chemiluminescent substrate (Thermo 34080) and imaged using Kodak X-OMAT 2000 Processor with BioMax light film (Carestream 178-8207).

Tissue was sampled from frozen hippocampi of a 73 year old male control (Braak=I, Thal=0) and an 86 year old male AD case (Braak=V, Thal=5). A second set of independent cases and controls was additionally investigated. For these, tissue was sampled from frozen mid-frontal cortices from three AD cases (#1 64 year old female [Braak=VI, Thal=5]; #2 60 year old female [Braak=VI, Thal=4]; #3 68 year old female [Braak VI; Thal 4]) and three controls (#4 75 year old female [Braak=I, Thal=0]; #5 78 year old male [Braak=II, Thal=1]; #6 96 year old female [Braak=II, Thal=3]). Raw western blot images can be found in **Extended Data Fig. 19-20**.

#### Statistical Considerations

To examine the association of neuropathologic markers with gene expression values, multivariable linear regression models were used to regress the observed gene expression values on digital pathology measures of AD pathology (tau [CP13, Ab39], Aβ [33.1.1]) and cellular diversity (activated microglia [CD68], endothelia [CD34], and astroglia [GFAP]), age at death, male sex, and presence of *APOE* ε4 risk allele. Models were fit, and the coefficients of partial determination were used to quantity the conditional association of each disease marker with the gene expression values. These coefficients of partial determination were combined across models for each gene in the form of a radar plot (**Fig. 4h**) to allow for visual comparison of how each gene may be differentially associated with neuropathologic markers.

All tests were two-sided and all p-values less than 0.05 were considered statistically significant. No correction for multiple testing has been applied to p-values to facilitate the application of other thresholds for statistical significance[99]. Continuous variables were summarized with median and range while categorical variables were summarized with frequency and percent. Mann-Whitney U tests with unadjusted post hoc comparisons of gene expression values across disease phenotypes were used to initially explore the data. All statistical analysis was performed in R Statistical Software (version 3.4.2; R Foundation for Statistical Computing, Vienna, Austria).
