## Extended data figures for "Leveraging selective hippocampal vulnerability among Alzheimer’s disease subtypes reveals a novel tau binding partner SERPINA5"

### Extended Data Fig. 1

| Characteristic | Controls (n=15) | AD neuropathologic subtypes (n=40) | | | AD specific p-value |
|---|---|---|---|---|---|
| | | HpSp AD (n=10) | Typical AD (n=20) | Limbic predominant AD (n=10) | |
| Males (% total of AD type) | 7/15 (47%) | 4/10 (40%) | 7/20 (35%) | 4/10 (40%) | 0.95 |
| *APOE* ε4, % | 1/15 (6.7%) | 3/10 (30%) | 8/20 (40%) | 4/10 (40%) | 0.85 |
| **Clinical findings** | | | | | |
| Age at onset, yr. | NA (NA, NA) | 66 (59, 69) | 74 (70, 80) | 79 (73, 80) | 0.006 |
| Disease duration, yr. | NA (NA, NA) | 8.7 (7.3, 8.9) | 7.7 (6.2, 10) | 9.0 (6.9, 10) | 0.76 |
| Final MMSE, points/yr. | 29 (28, 30) | 13 (6.5, 16) | 14 (12, 19) | 14 (7.0, 21) | 0.68 |
| **Postmortem findings** | | | | | |
| Age at death, yr. | 84 (80, 90) | 73 (68, 76) | 81 (78, 88) | 87 (82, 89) | 0.003 |
| Braak tangle stage | II (I, III) | VI (V, VI) | VI (V, VI) | VI (V, VI) | 0.95 |
| Thal amyloid phase | 0 (0, 1) | 5 (5, 5) | 5 (4, 5) | 5 (5, 5) | 0.38 |
| Early tangle marker (CP13), % | 0.22 (0.10, 1.5) | 12 (8.2, 14) | 12 (9.9, 17) | 16 (13, 26) | 0.11 |
| Mature tangle marker (Ab39), % | 0.040 (0.030, 0.080) | 0.22 (0.19, 0.43) | 0.27 (0.20, 0.37) | 0.53 (0.44, 0.76) | 0.018 |
| Pan-Aβ marker (33.1.1), % | 0.20 (0.080, 0.33) | 0.56 (0.38, 0.81) | 0.56 (0.43, 1.1) | 0.50 (0.47, 0.60) | 0.94 |
| Astroglial marker (GFAP), % | 29 (20, 36) | 34 (22, 39) | 36 (31, 47) | 36 (32, 50) | 0.34 |
| Activated microglial marker (CD68), % | 0.10 (0.060, 0.20) | 0.24 (0.17, 0.37) | 0.32 (0.22, 0.38) | 0.51 (0.37, 0.63) | 0.095 |
| Endothelial cell marker (CD34), % | 0.72 (0.50, 0.81) | 0.52 (0.34, 0.64) | 0.65 (0.37, 0.87) | 0.87 (0.72, 0.93) | 0.041 |

### Extended Data Fig. 2

| Corresponding figure | Description | Control vs HpSp AD | Control vs Typical AD | Control vs Limbic AD | HpSp AD vs Typical AD | HpSp AD vs Limbic AD | Typical AD vs Limbic AD |
|---|---|---|---|---|---|---|---|
| Fig. 3f | *SIRT1* levels from RNA-Seq | 0.31 | **0.034** | **0.0029** | 0.38 | **0.015** | 0.12 |
| Fig. 3g | *PSEN2* levels from RNA-Seq | 0.37 | **0.011** | **0.004** | 0.19 | 0.11 | 0.68 |
| Fig. 5a | *SERPINA5* levels from RNA-Seq | **0.001** | **<0.001** | **0.004** | 0.56 | 0.58 | 0.81 |
| Fig. 5b | *SERPINA5* levels from NanoString | **<0.001** | **<0.001** | **<0.001** | 0.9 | **0.023** | **0.011** |
| Fig. 5d | CA1 *SERPINA5* burden | **<0.001** | **<0.001** | **<0.001** | **0.026** | 0.076 | 0.99 |
| Fig. 5d | Subiculum *SERPINA5* burden | **<0.001** | **<0.001** | **<0.001** | 0.096 | **0.023** | 0.64 |
| Fig. 5d | Superior temporal *SERPINA5* burden | **<0.001** | **<0.001** | **<0.001** | 0.9 | **0.006** | 0.14 |
| Fig. 5d | Inferior temporal *SERPINA5* burden | **<0.001** | **<0.001** | **<0.001** | 0.24 | **<0.001** | **0.002** |
| Fig. 5d | Mid-frontal *SERPINA5* burden | **<0.001** | **<0.001** | **<0.001** | 0.056 | **<0.001** | **<0.001** |

### Extended Data Fig. 3

| Gene selection | Phenotype derived | Genes | Transcript level associations with neuropathologic measures (false discovery rate<0.25, p-value<0.05) | | | | | | | | | | Genes found to robustly associate in both analyses (All four groups and Representative phenotype) | | | | |
|---|---|---|---|---|---|---|---|---|---|---|---|---|---|---|---|---|---|
| | | | All four groups | | | | | Representative phenotype | | | | | | | | | |
| | | | Tau | | | Amyloid-β | | Tau | | | Amyloid-β | | Tau | | | Amyloid-β | |
| | | | Braak | CP13 | Ab39 | Thal | 33.1.1 | Braak | CP13 | Ab39 | Thal | 33.1.1 | Braak | CP13 | Ab39 | Thal | 33.1.1 |
| Step 1: AD Literature | n/a | ***PSEN2*** | | | | | | | | | | | | | | | |
| | n/a | ***SIRT1*** | | | | | | | | | | | | | | | |
| Step 2: Bioinformatic Prioritization | Representative | ***ALOX15B*** | | | | | | | | | | | | | | | |
| | Representative | ***ANGPT2*** | | | | | | | | | | | | | | | |
| | Representative | ***CXCL1*** | | | | | | | | | | | | | | | |
| | Representative | ***DNAAF1*** | | | | | | | | | | | | | | | |
| | Representative | ***DNAI1*** | | | | | | | | | | | | | | | |
| | Extreme | ***DYDC2*** | | | | | | | | | | | | | | | |
| | Representative | ***LRRC48*** | | | | | | | | | | | | | | | |
| | Both | ***MAPK15*** | | | | | | | | | | | | | | | |
| | Representative | ***OR7A5*** | | | | | | | | | | | | | | | |
| | Representative | ***PYDC1*** | | | | | | | | | | | | | | | |
| | Extreme | ***RBP1*** | | | | | | | | | | | | | | | |
| | Representative | ***RP11-81K2.1*** | | | | | | | | | | | | | | | |
| | Representative | ***SERPINA5*** | | | | | | | | | | | | | | | |
| | Representative | ***TAC1*** | | | | | | | | | | | | | | | |
| Step 3: Process Networks | Representative | ***ATR*** | | | | | | | | | | | | | | | |
| | Representative | ***BCL2*** | | | | | | | | | | | | | | | |
| | Extreme | ***CAV1*** | | | | | | | | | | | | | | | |
| | Representative | ***CDKN2C*** | | | | | | | | | | | | | | | |
| | Representative | ***CNOT8*** | | | | | | | | | | | | | | | |
| | Representative | ***CTCF*** | | | | | | | | | | | | | | | |
| | Representative | ***CXCL1*** | | | | | | | | | | | | | | | |
| | Representative | ***DAXX*** | | | | | | | | | | | | | | | |
| | Extreme | ***DST*** | | | | | | | | | | | | | | | |
| | Representative | ***EML4*** | | | | | | | | | | | | | | | |
| | Extreme | ***ERBB2IP*** | | | | | | | | | | | | | | | |
| | Representative | ***FEM1B*** | | | | | | | | | | | | | | | |
| | Representative | ***FOXO4*** | | | | | | | | | | | | | | | |
| | Extreme | ***IFITM2*** | | | | | | | | | | | | | | | |
| | Representative | ***INSR*** | | | | | | | | | | | | | | | |
| | Representative | ***IRS2*** | | | | | | | | | | | | | | | |
| | Representative | ***LCOR*** | | | | | | | | | | | | | | | |
| | Extreme | ***MAGED1*** | | | | | | | | | | | | | | | |
| | Representative | ***MAX*** | | | | | | | | | | | | | | | |
| | Representative | ***NCOA1*** | | | | | | | | | | | | | | | |
| | Representative | ***NOS1*** | | | | | | | | | | | | | | | |
| | Representative | ***PPM1D*** | | | | | | | | | | | | | | | |
| | Representative | ***RAD52*** | | | | | | | | | | | | | | | |
| | Representative | ***RBBP7*** | | | | | | | | | | | | | | | |
| | Representative | ***RELA*** | | | | | | | | | | | | | | | |
| | Representative | ***RYBP*** | | | | | | | | | | | | | | | |
| | Representative | ***SIPA1*** | | | | | | | | | | | | | | | |
| | Representative | ***SLC38A2*** | | | | | | | | | | | | | | | |
| | Representative | ***SUN2*** | | | | | | | | | | | | | | | |
| **Total Genes Associated*** | | | 30 | 0 | 35 | 36 | 0 | 13 | 33 | 35 | 23 | 0 | 13 | 0 | 35 | 23 | 0 |

### Extended Data Fig. 4

| Characteristic | Controls (n=32) | AD neuropathologic subtypes (n=150) | | | AD specific p-value |
|---|---|---|---|---|---|
| | | HpSp AD (n=36) | Typical AD (n=79) | Limbic predominant AD (n=35) | |
| Males (% total of AD type) | 15/32 (47%) | 14/36 (39%) | 26/79 (33%) | 16/35 (46%) | 0.42 |
| *APOE* ε4, % | 5/32 (16%) | 14/36 (39%) | 40/79 (51%) | 23/35 (66%) | 0.076 |
| **Clinical findings** | | | | | |
| Age at onset, yr. | NA (NA, NA) | 66 (60, 72) | 73 (67, 79) | 79 (72, 82) | <0.001 |
| Disease duration, yr. | NA (NA, NA) | 8.4 (6.6, 9.3) | 8.3 (6.3, 10) | 9.4 (6.9, 11) | 0.36 |
| Final MMSE, points/yr. | 29 (27, 29) | 8 (1.5, 12) | 14 (9.3, 22) | 16 (8, 22) | 0.15 |
| **Postmortem findings** | | | | | |
| Age at death, yr. | 87 (80, 91) | 72 (68, 80) | 82 (74, 88) | 87 (82, 93) | <0.001 |
| Braak tangle stage | II (II, III) | VI (V, VI) | VI (V, VI) | VI (V, VI) | 0.26 |
| Thal amyloid phase | 0 (0, 2) | 5 (5, 5) | 5 (5, 5) | 5 (4, 5) | 0.097 |
| Early tangle marker (CP13), % | 0.4 (0.1, 1.7) | 11 (7.6, 16) | 15 (11, 18) | 15 (11, 19) | 0.12 |
| Mature tangle marker (Ab39), % | 0.06 (0.04, 0.09) | 0.24 (0.11, 0.43) | 0.33 (0.21, 0.64) | 0.54 (0.39, 0.77) | <0.001 |
| Pan-Aβ marker (33.1.1), % | 0.12 (0.07, 0.21) | 0.38 (0.24, 0.62) | 0.41 (0.31, 0.59) | 0.49 (0.32, 0.64) | 0.58 |
| Astrogliosis marker (GFAP), % | 31 (23, 37) | 33 (23, 40) | 43 (34, 52) | 53 (43, 59) | <0.001 |
| Activated microglial marker (CD68), % | 0.13 (0.08, 0.21) | 0.23 (0.17, 0.34) | 0.30 (0.22, 0.41) | 0.38 (0.30, 0.54) | 0.0020 |
| Endothelial cell marker (CD34), % | 0.73 (0.58, 0.83) | 0.47 (0.37, 0.78) | 0.67 (0.49, 0.97) | 0.90 (0.67, 0.99) | <0.001 |

### Extended Data Fig. 5

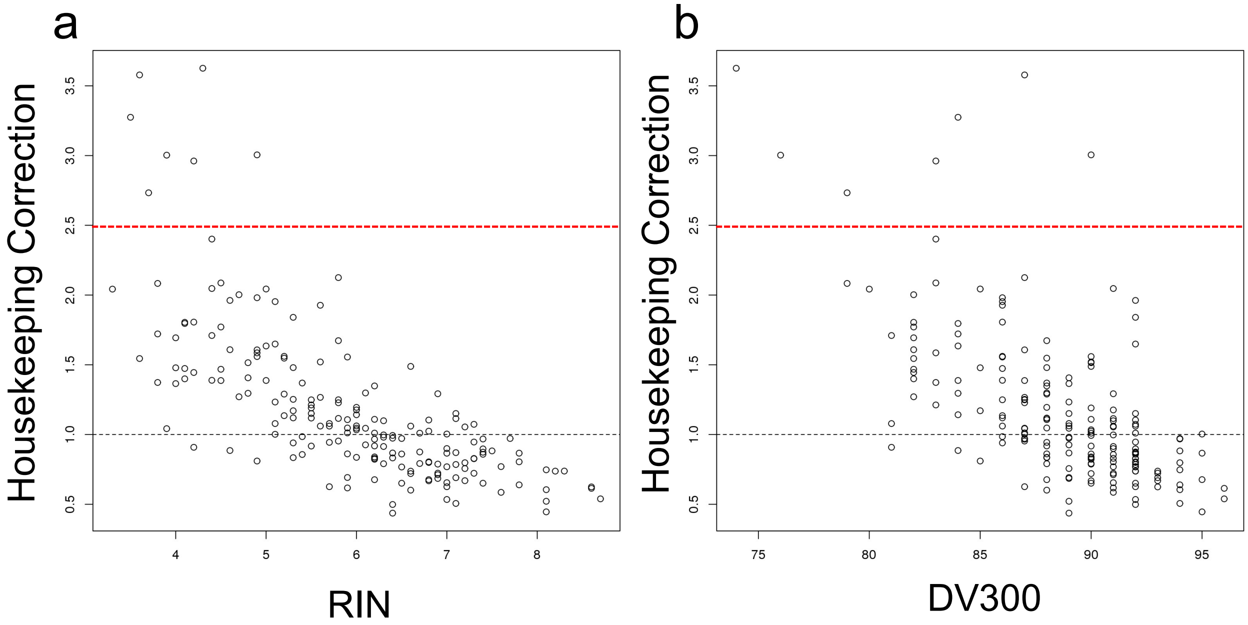

### Extended Data Fig. 6

| Gene | Pearson Correlation | R-squared | p-value |
|---|---|---|---|
| *SERPINA5* | 0.899 | 0.81 | <0.001 |
| *RYBP* | 0.83 | 0.69 | <0.001 |
| *SLC38A2* | 0.99 | 0.97 | <0.001 |
| *FEM1B* | 0.83 | 0.68 | <0.001 |
| *PYDC1* | 0.74 | 0.55 | <0.001 |

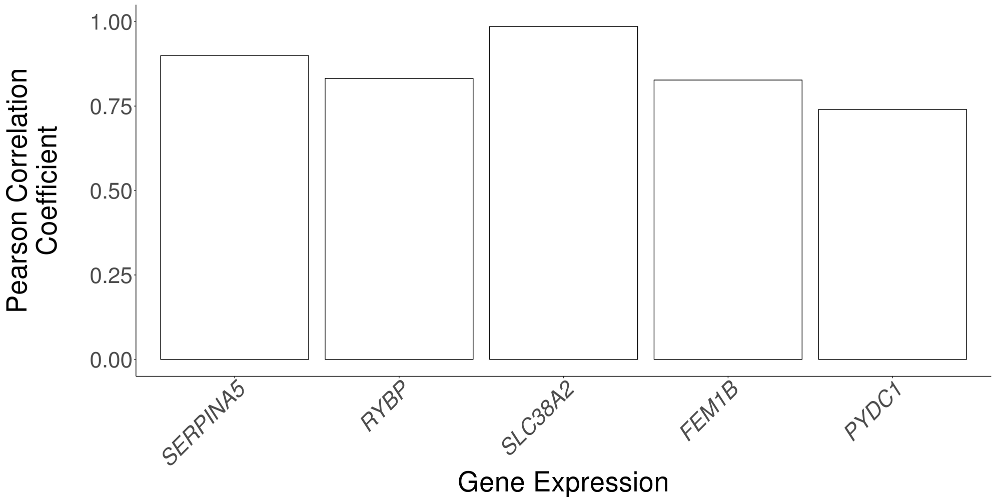

Extended Data Fig. 7

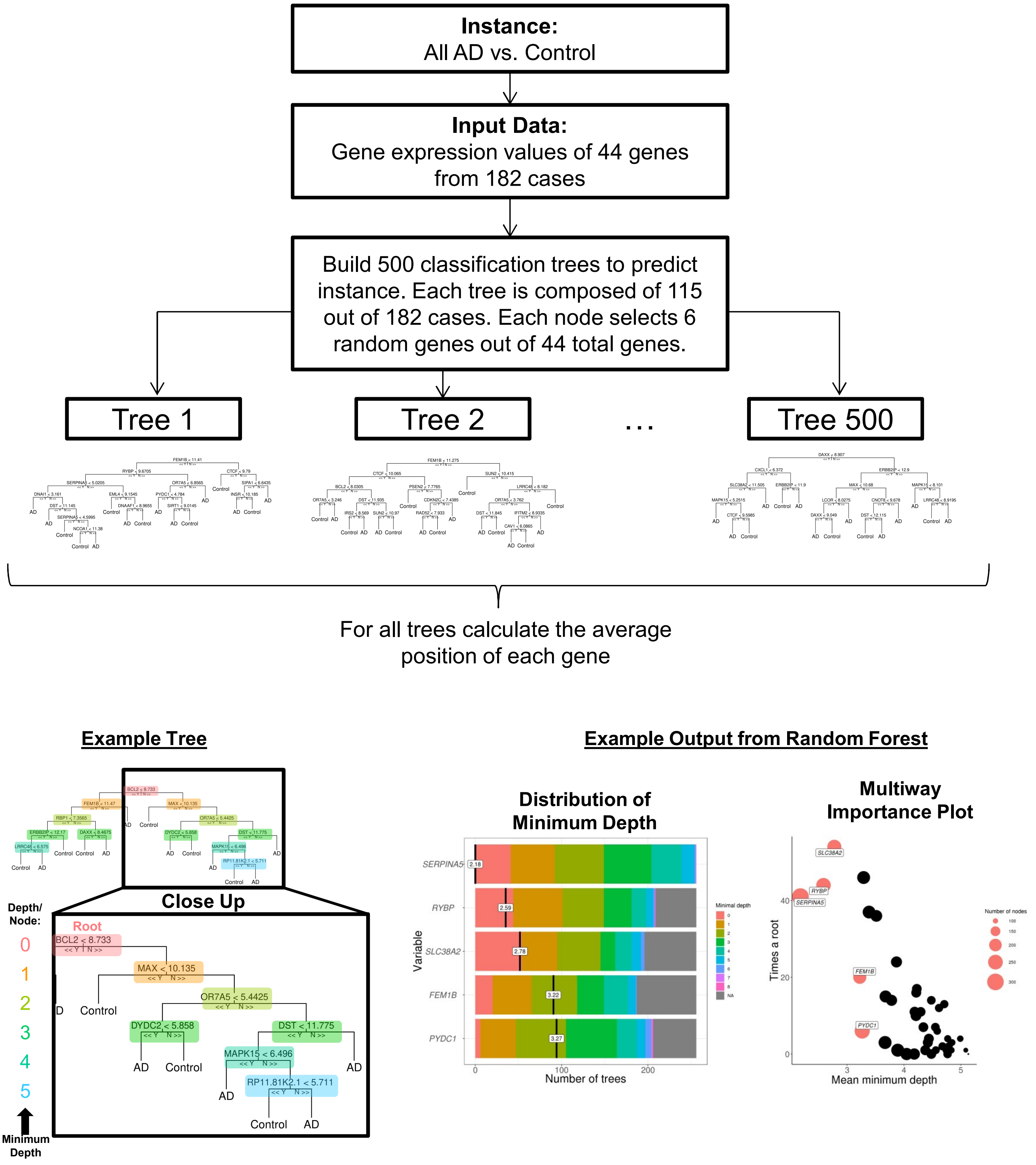

### Extended Data Fig. 8

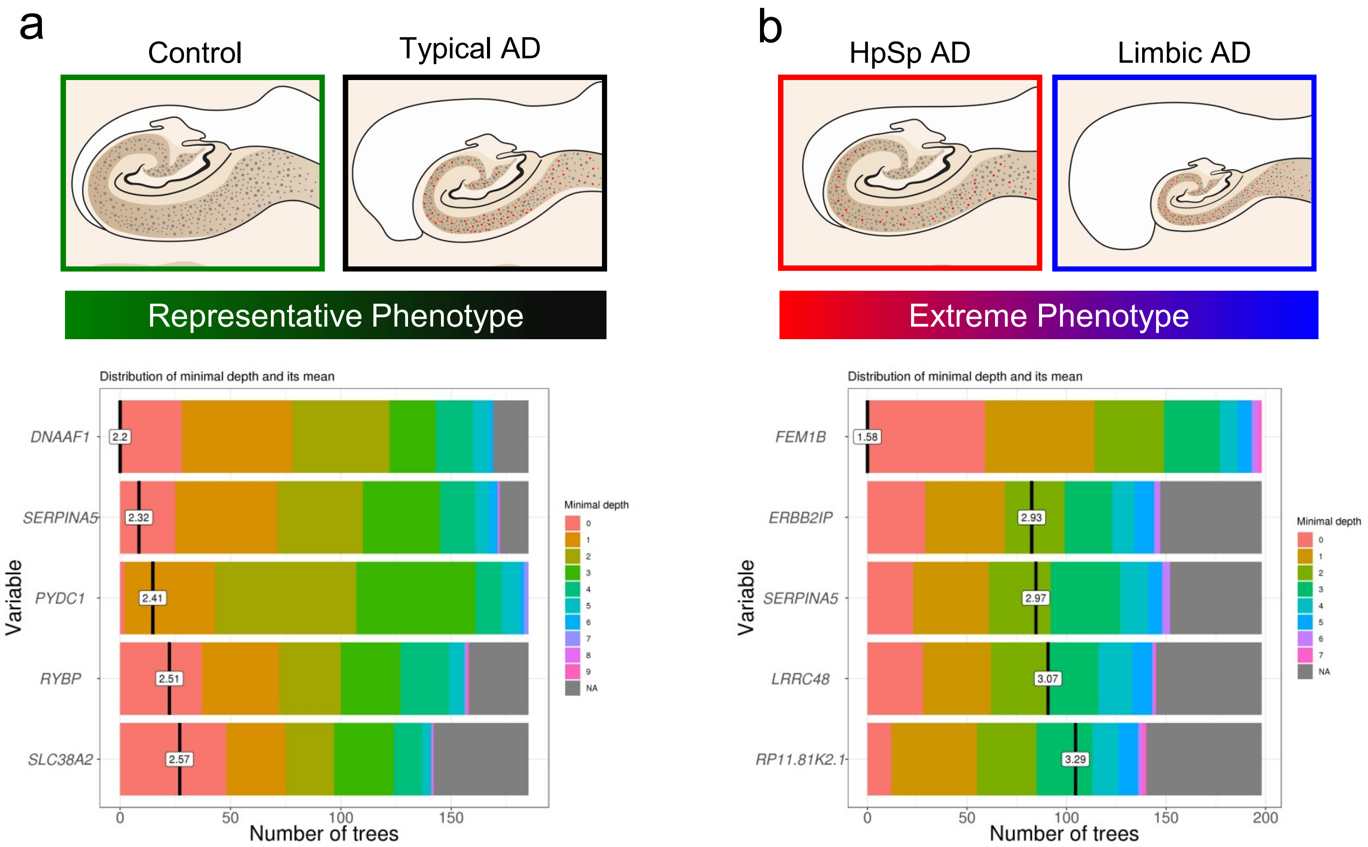

### Extended Data Fig. 9

| **Variable (unit change)** | **OR (95% CI)** | **p-value** |
|---|---|---|
| Sex (male) | 0.67 (0.24, 1.9) | 0.45 |
| Age (15 years) | 0.57 (0.23, 1.4) | 0.23 |
| *SERPINA5* gene expression (2.0) | 2.3 (1.3, 4.2) | 0.0040 |
| *SLC38A2* gene expression (1.2) | 7.6 (1.5, 39) | 0.016 |
| *FEM1B* gene expression (0.38) | 2.4 (0.82, 7.4) | 0.11 |
| *PYDC1* gene expression (1.4) | 0.49 (0.25, 0.97) | 0.040 |
| *RYBP* gene expression (0.51) | 0.77 (0.26, 2.3) | 0.65 |

### Extended Data Fig. 10

a **How to use a nomogram**

1. Plot sex, age and gene expression values (■)
2. Add up total points (↑)
3. Plot total points (■)
4. Obtain predicted value (↓■)

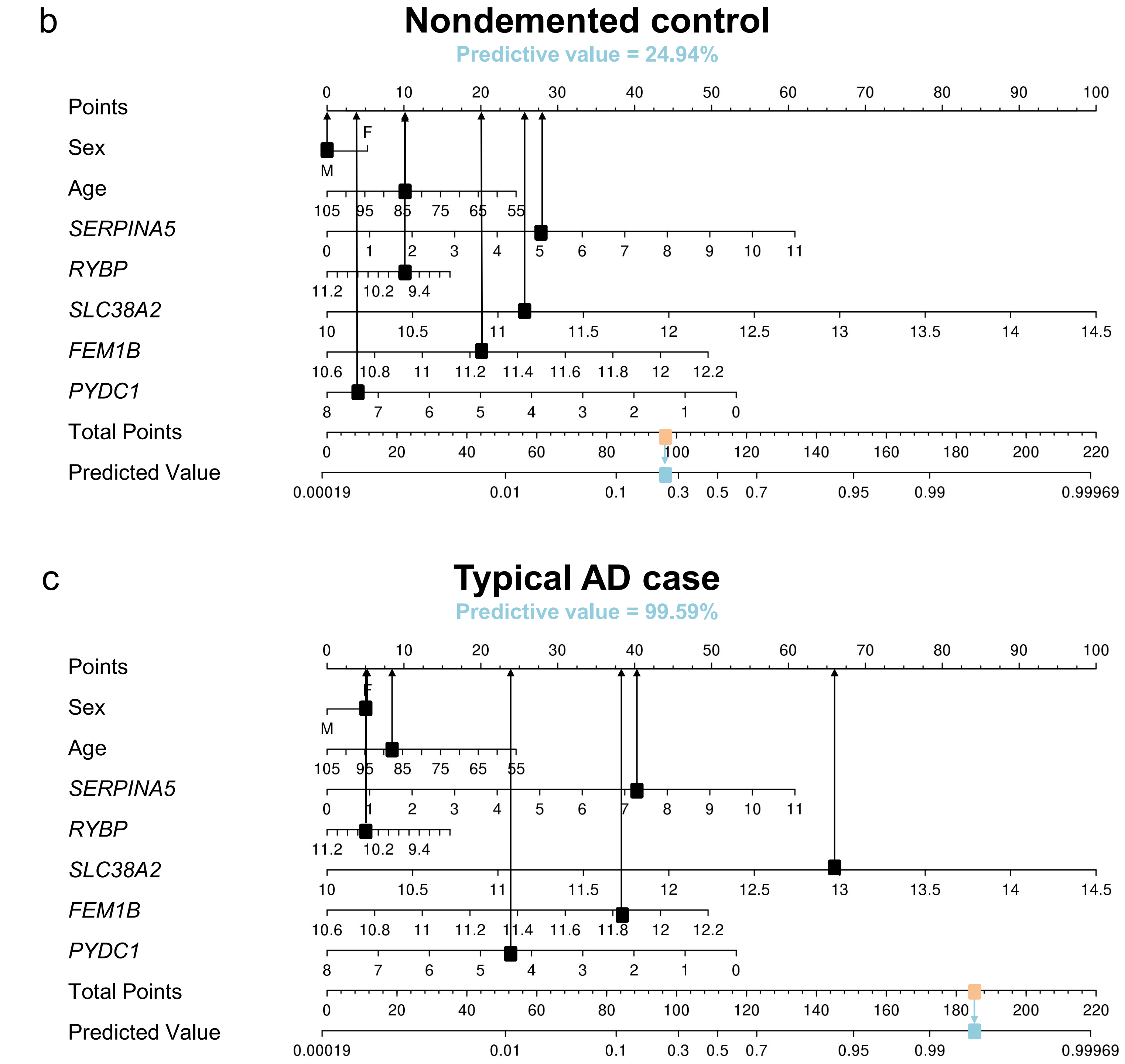

### Extended Data Fig. 11

| Markers and covariates | *SERPINA5* | | *RYBP* | | *SLC38A2* | | *FEM1B* | | *PYDC1* | |
|---|---|---|---|---|---|---|---|---|---|---|
| | **Coeff.** | **P-value** | **Coeff.** | **P-value** | **Coeff.** | **P-value** | **Coeff.** | **P-value** | **Coeff.** | **P-value** |
| Early tangle | 0.002 | 0.62 | 0.014 | 0.14 | **0.040** | **0.011** | **0.076** | **0.000** | **0.030** | **0.028** |
| Advanced tangle | **0.041** | **0.010** | 0.015 | 0.13 | **0.031** | **0.026** | 0.00 | 0.88 | 0.007 | 0.29 |
| Pan-Aβ | 0.002 | 0.58 | 0.002 | 0.63 | 0.001 | 0.74 | 0.016 | 0.11 | 0.000 | 0.99 |
| Microglial | 0.020 | 0.075 | 0.024 | 0.052 | 0.024 | 0.052 | 0.019 | 0.087 | 0.000 | 0.88 |
| Endothelial | 0.014 | 0.14 | 0.017 | 0.097 | 0.012 | 0.17 | 0.008 | 0.25 | 0.000 | 0.79 |
| Astroglial | 0.004 | 0.40 | 0.002 | 0.60 | 0.002 | 0.58 | 0.039 | 0.013 | 0.000 | 0.84 |
| Age at death | 0.004 | 0.40 | **0.082** | **0.00** | 0.022 | 0.060 | **0.054** | **0.003** | 0.024 | 0.052 |
| Sex (Male) | 0.014 | 0.13 | 0.00 | 0.90 | 0.00 | 0.88 | 0.001 | 0.67 | 0.004 | 0.44 |
| *APOE ε*4 status | 0.00 | 0.99 | 0.017 | 0.098 | 0.017 | 0.10 | 0.001 | 0.74 | 0.000 | 0.81 |

Extended Data Fig. 12

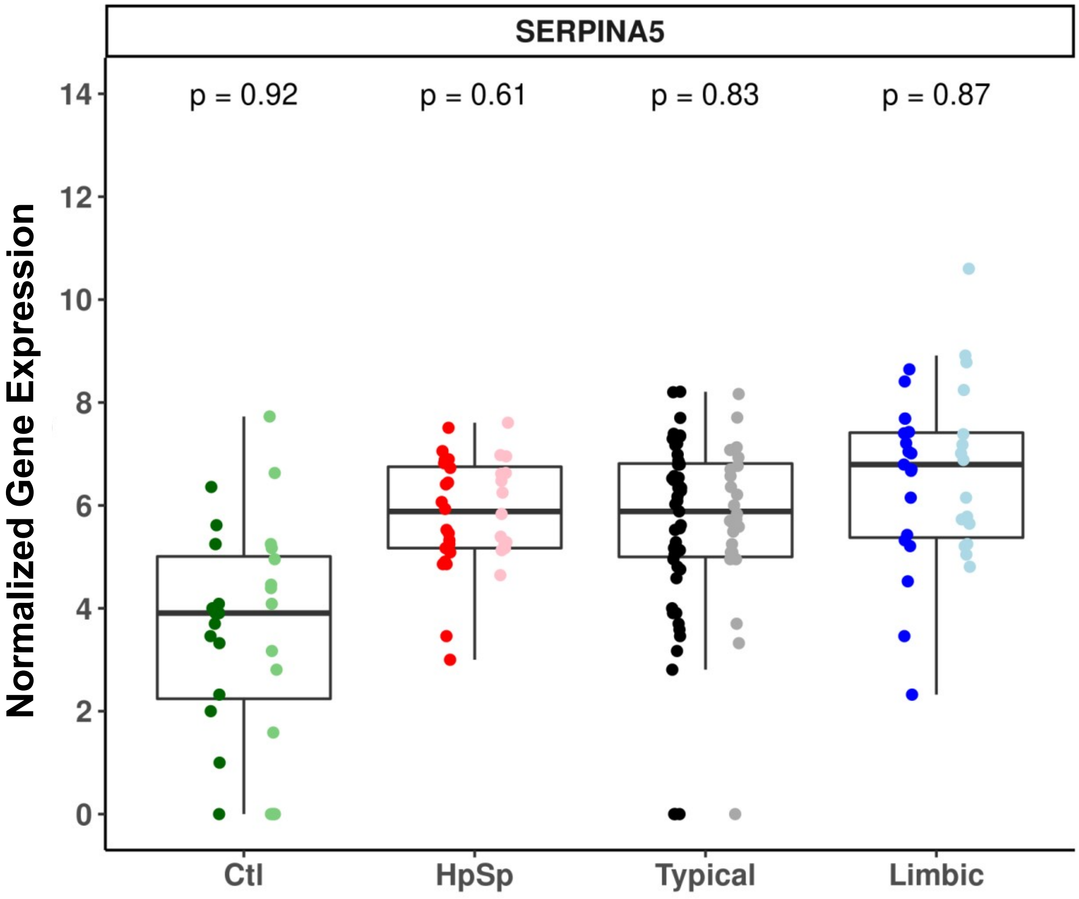


Note: Each jitter plot overlay is stratified by sex with females on the left (darker color) and males on the right (lighter color)

Extended Data Fig. 13

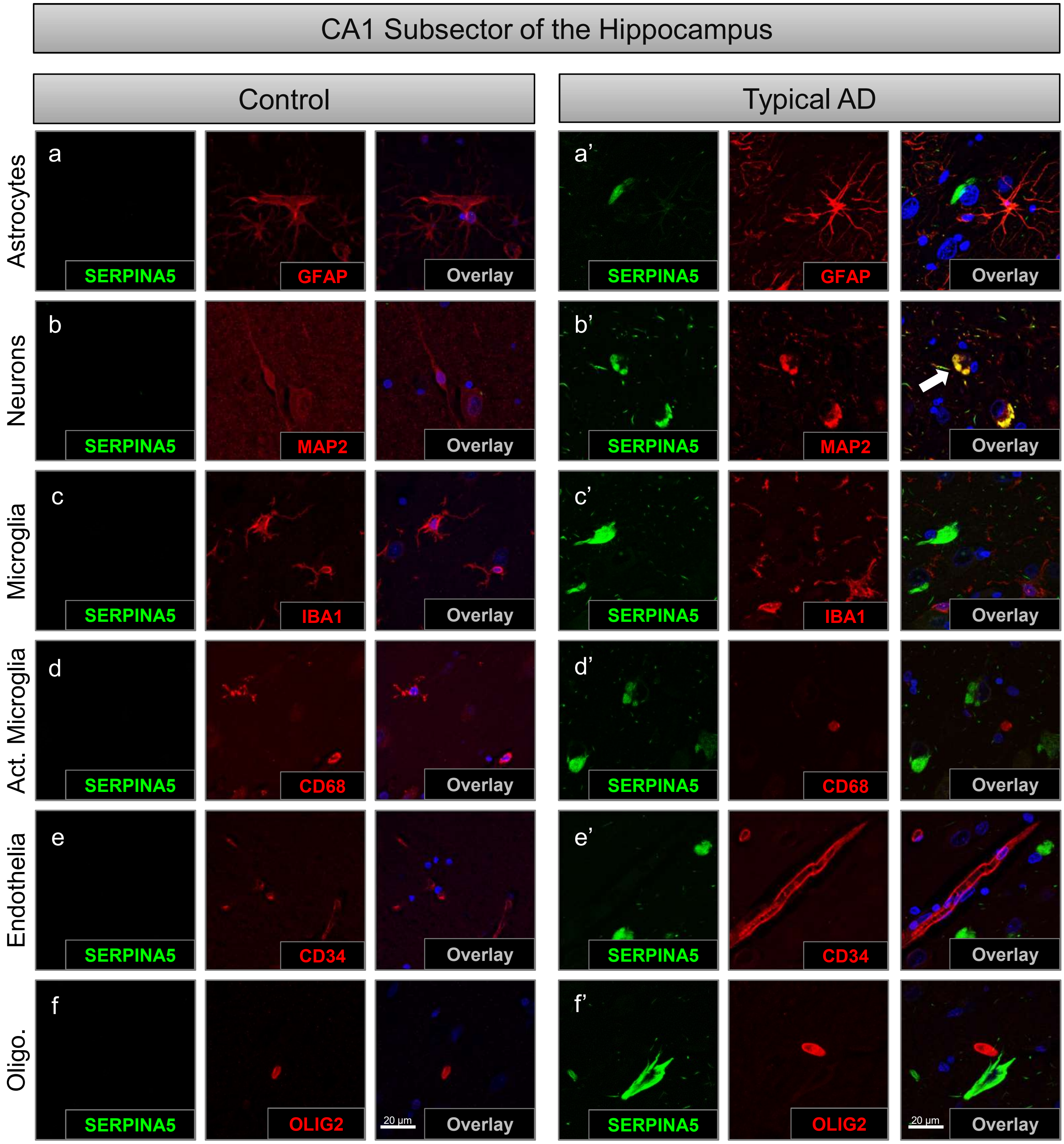

Extended Data Fig. 14

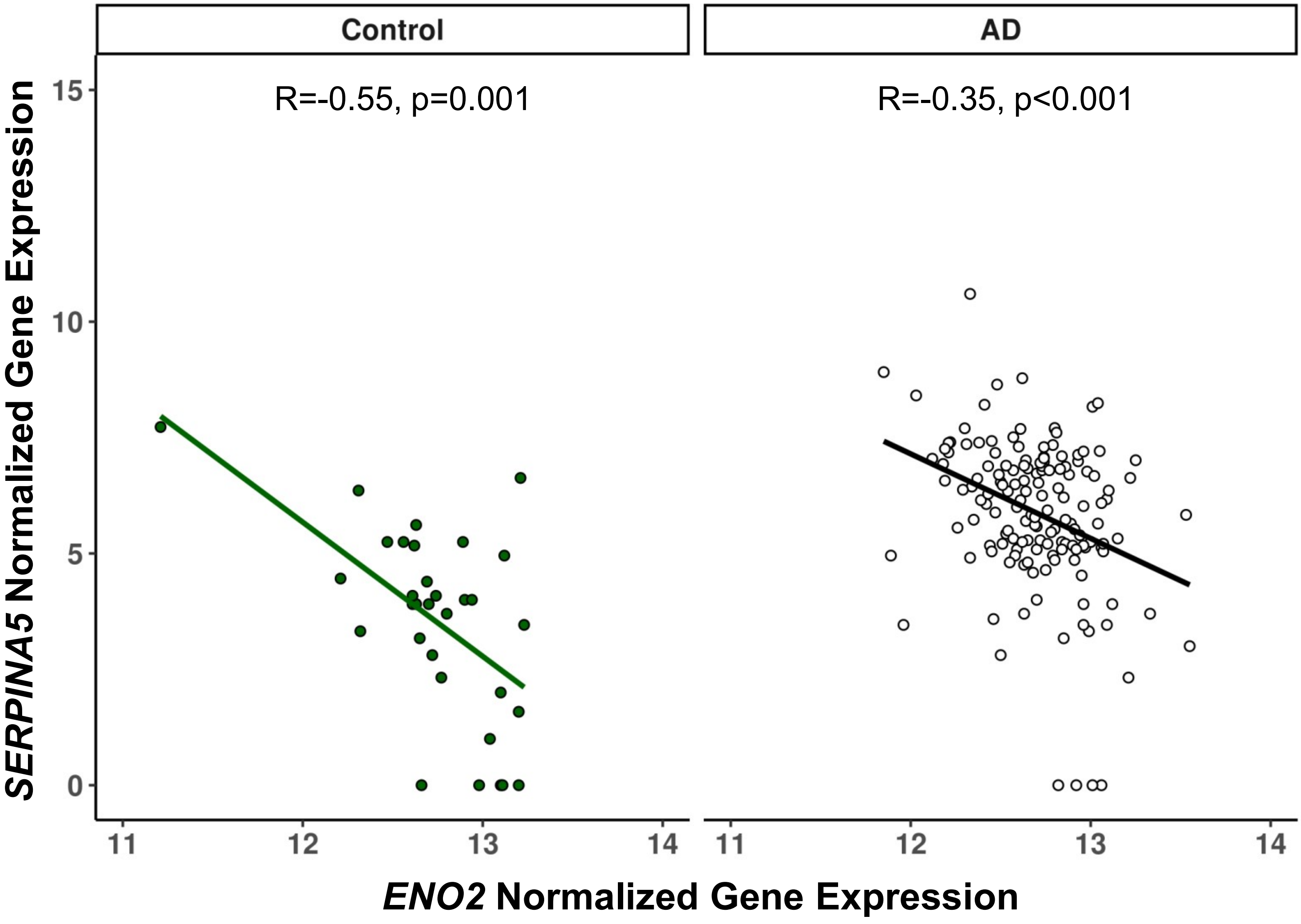

### Extended Data Fig. 15

| Characteristic | Controls (n=10) | AD neuropathologic subtypes (n=40) | | | AD specific p-value |
|---|---|---|---|---|---|
| | | HpSp AD (n=20) | Typical AD (n=20) | Limbic predominant AD (n=20) | |
| Males (% total of AD type) | 5/10 (50%) | 15/20 (75%) | 11/20 (55%) | 3/20 (15%) | < 0.001 |
| *APOE* ε4*, % | 0/1 (0%) | 10/17 (59%) | 13/19 (68%) | 6/9 (67%) | 0.83 |
| **Clinical findings** | | | | | |
| Age at onset, yr. | NA (NA, NA) | 65 (58, 68) | 72 (65, 76) | 81 (78, 87) | < 0.001 |
| Disease duration, yr. | NA (NA, NA) | 7.9 (6.3, 10) | 10 (7.3, 15) | 6.8 (6, 8.6) | 0.18 |
| **Postmortem findings** | | | | | |
| Age at death, yr. | 76 (60, 81) | 71 (68, 74) | 82 (76, 85) | 87 (84, 91) | < 0.001 |
| Braak tangle stage | I (I, II) | VI (V, VI) | VI (V, VI) | V (IV,VI) | 0.011 |
| Thal amyloid phase | 1 (0, 2) | 5 (5, 5) | 5 (5, 5) | 5 (5, 5) | 0.87 |
| Average hippocampal SERPINA5, % | 0.010 (0.010, 0.020) | 2.6 (1.8, 3.9) | 4.0 (2.4, 8.4) | 5.3 (2.0, 9.8) | 0.011 |
| CA1, % | 0.010 (0.010, 0.020) | 1.0 (0.77, 1.4) | 2.0 (0.94, 2.8) | 1.8 (0.78, 3.8) | 0.015 |
| Subiculum, % | 0.020 (0.010, 0.020) | 1.6 (0.76, 2.4) | 2.2 (1.4, 4.9) | 3.5 (1.2, 5.6) | 0.014 |
| Average cortical SERPINA5, % | 0.010 (0.010, 0.020) | 5.8 (3.9, 7.6) | 5.4 (1.4, 7.6) | 1.7 (0.95, 2.8) | <0.001 |
| Superior temporal, % | 0.010 (0.010, 0.020) | 1.2 (0.85, 1.7) | 1.4 (0.44, 2.8) | 0.71 (0.43, 1.1) | 0.075 |
| Inferior parietal, % | 0.010 (0.010, 0.020) | 1.8 (1.2, 2.6) | 1.4 (0.54, 3.3) | 0.44 (0.23, 0.67) | <0.001 |
| Mid-frontal, % | 0.010 (0.010, 0.020) | 2.2 (1.6, 3.2) | 1.8 (0.53, 2.2) | 0.19 (0.12, 0.43) | <0.001 |

### Extended Data Fig. 16

| AD Subtype | Total reads | Used reads | Mapped reads | Mapped reads (Genome) | Mapped reads (Junction) | Gene count | Exon count |
|---|---|---|---|---|---|---|---|
| Ctl-1 | 107,651,934 | 107,568,111 | 93,194,746 (86.6) | 83,514,090 (77.6) | 9,680,656 (9.0) | 33,353,018 (31.0) | 173,910,437 (161.5) |
| Ctl-2 | 143,957,388 | 143,812,328 | 123,167,238 (85.6) | 111,860,651 (77.7) | 11,306,587 (7.9) | 43,690,722 (30.3) | 236,111,216 (164.0) |
| Ctl-3 | 179,380,730 | 179,357,440 | 147,567,698 (82.3) | 129,896,840 (72.4) | 17,670,858 (9.9) | 61,174,456 (34.1) | 330,677,695 (184.3) |
| Ctl-4 | 118,329,800 | 118,199,155 | 98,321,008 (83.1) | 86,910,578 (73.4) | 11,410,430 (9.6) | 38,537,231 (32.6) | 209,250,869 (176.8) |
| Ctl-5 | 134,746,118 | 134,617,790 | 116,033,251 (86.1) | 98,736,415 (73.3) | 17,296,836 (12.8) | 53,015,332 (39.3) | 293,307,193 (217.7) |
| Ctl-6 | 118,919,334 | 118,857,614 | 103,343,443 (86.9) | 89,839,574 (75.5) | 13,503,869 (11.4) | 43,966,262 (37.0) | 240,414,294 (202.2) |
| Ctl-7 | 102,808,398 | 102,728,580 | 88,127,185 (85.7) | 76,617,237 (74.5) | 11,509,948 (11.2) | 37,153,641 (36.1) | 209,289,438 (203.6) |
| Ctl-8 | 153,409,250 | 153,254,804 | 128,699,480 (83.9) | 114,521,058 (74.7) | 14,178,422 (9.2) | 48,480,606 (31.6) | 264,765,534 (172.6) |
| Ctl-9 | 112,510,692 | 112,408,836 | 93,514,202 (83.1) | 83,352,847 (74.1) | 10,161,355 (9.0) | 37,432,246 (33.3) | 209,566,548 (186.3) |
| Ctl-10 | 102,883,862 | 102,801,693 | 86,186,616 (83.8) | 76,974,145 (74.8) | 9,212,471 (9.0) | 34,656,660 (33.7) | 188,725,481 (183.4) |
| Ctl-11 | 138,409,450 | 138,268,143 | 114,903,154 (83.0) | 104,115,662 (75.2) | 10,787,492 (7.8) | 42,052,850 (30.4) | 224,181,633 (162.0) |
| Ctl-12 | 137,351,268 | 137,208,841 | 117,197,326 (85.3) | 105,333,157 (76.7) | 11,864,169 (8.6) | 42,096,302 (30.6) | 230,046,259 (167.5) |
| Ctl-13 | 125,903,118 | 125,780,042 | 107,610,184 (85.5) | 90,092,659 (71.6) | 17,517,525 (13.9) | 54,993,809 (43.7) | 311,179,897 (247.2) |
| Ctl-14 | 165,767,984 | 165,747,538 | 140,771,037 (84.9) | 123,432,263 (74.5) | 17,338,774 (10.5) | 57,999,533 (35.0) | 319,470,580 (192.7) |
| Ctl-15 | 127,584,888 | 127,458,173 | 108,436,878 (85.0) | 94,494,689 (74.1) | 13,942,189 (10.9) | 45,757,687 (35.9) | 252,590,533 (198.0) |
| HpSp-1 | 125,276,906 | 125,170,419 | 106,938,562 (85.4) | 98,670,718 (78.8) | 8,267,844 (6.6) | 31,258,536 (25.0) | 162,297,018 (129.6) |
| HpSp-2 | 112,391,398 | 112,298,101 | 96,120,757 (85.5) | 87,315,274 (77.7) | 8,805,483 (7.8) | 34,170,760 (30.4) | 182,357,957 (162.3) |
| HpSp-3 | 124,563,828 | 124,462,427 | 106,689,763 (85.7) | 94,358,412 (75.8) | 12,331,351 (9.9) | 44,222,969 (35.5) | 237,704,396 (190.8) |
| HpSp-4 | 257,886,536 | 257,843,755 | 223,556,514 (86.7) | 197,410,263 (76.5) | 26,146,251 (10.1) | 97,530,807 (37.8) | 548,977,311 (212.9) |
| HpSp-5 | 240,891,618 | 240,837,051 | 204,623,466 (84.9) | 180,740,549 (75.0) | 23,882,917 (9.9) | 88,254,170 (36.6) | 479,412,236 (199.0) |
| HpSp-6 | 108,879,028 | 108,788,404 | 92,934,112 (85.4) | 79,679,966 (73.2) | 13,254,146 (12.2) | 39,141,681 (35.9) | 214,916,975 (197.4) |
| HpSp-7 | 135,137,536 | 135,003,369 | 115,152,555 (85.2) | 101,068,299 (74.8) | 14,084,256 (10.4) | 45,468,489 (33.6) | 239,973,913 (177.6) |
| HpSp-8 | 102,379,238 | 102,298,930 | 89,748,321 (87.7) | 80,618,042 (78.7) | 9,130,279 (8.9) | 35,746,929 (34.9) | 191,239,060 (186.8) |
| HpSp-9 | 103,100,460 | 103,016,548 | 90,481,939 (87.8) | 81,212,317 (78.8) | 9,269,622 (9.0) | 33,389,408 (32.4) | 177,644,705 (172.3) |
| HpSp-10 | 111,566,254 | 111,459,962 | 97,426,140 (87.3) | 85,344,209 (76.5) | 12,081,931 (10.8) | 41,897,472 (37.6) | 229,765,923 (205.9) |
| Typical-1 | 97,194,370 | 97,119,394 | 83,681,667 (86.1) | 73,699,151 (75.8) | 9,982,516 (10.3) | 36,072,349 (37.1) | 193,412,105 (199.0) |
| Typical-2 | 340,499,070 | 340,422,280 | 284,816,399 (83.6) | 248,762,075 (73.1) | 36,054,324 (10.6) | 119,359,928 (35.1) | 633,471,756 (186.0) |
| Typical-3 | 84,807,392 | 84,735,639 | 72,524,774 (85.5) | 64,581,000 (76.2) | 7,943,774 (9.4) | 30,375,552 (35.8) | 165,606,914 (195.3) |
| Typical-4 | 118,105,530 | 117,990,305 | 101,555,777 (86.0) | 91,640,778 (77.6) | 9,914,999 (8.4) | 39,511,999 (33.5) | 205,378,512 (173.9) |
| Typical-5 | 146,395,500 | 146,270,388 | 124,619,457 (85.1) | 113,148,447 (77.3) | 11,471,010 (7.8) | 45,972,822 (31.4) | 241,880,059 (165.2) |
| Typical-6 | 127,957,452 | 127,826,202 | 107,764,096 (84.2) | 95,164,934 (74.4) | 12,599,162 (9.8) | 40,650,935 (31.8) | 219,897,728 (171.9) |
| Typical-7 | 140,652,042 | 140,510,428 | 118,797,157 (84.5) | 103,874,665 (73.9) | 14,922,492 (10.6) | 51,881,008 (36.9) | 280,995,043 (199.8) |
| Typical-8 | 169,673,950 | 169,651,345 | 135,795,507 (80.0) | 121,626,439 (71.7) | 14,169,068 (8.4) | 53,375,589 (31.5) | 285,082,193 (168.0) |
| Typical-9 | 261,480,608 | 261,434,148 | 226,628,485 (86.7) | 199,107,889 (76.1) | 27,520,596 (10.5) | 99,517,980 (38.1) | 553,777,854 (211.8) |
| Typical-10 | 148,154,652 | 148,017,561 | 124,021,870 (83.7) | 112,545,054 (76.0) | 11,476,816 (7.7) | 42,183,333 (28.5) | 220,340,383 (148.7) |
| Typical-11 | 112,105,860 | 112,016,935 | 94,369,236 (84.2) | 84,816,813 (75.7) | 9,552,423 (8.5) | 36,967,227 (33.0) | 198,202,069 (176.8) |
| Typical-12 | 104,536,288 | 104,450,587 | 89,427,140 (85.5) | 80,053,842 (76.6) | 9,373,298 (9.0) | 32,423,808 (31.0) | 173,988,951 (166.4) |
| Typical-13 | 112,087,790 | 111,979,772 | 96,506,791 (86.1) | 85,997,307 (76.7) | 10,509,484 (9.4) | 37,243,917 (33.2) | 203,086,222 (181.2) |
| Typical-14 | 154,244,518 | 154,165,681 | 130,499,592 (84.6) | 114,971,350 (74.5) | 15,528,242 (10.1) | 55,799,738 (36.2) | 308,004,109 (199.7) |
| Typical-15 | 113,086,410 | 112,994,111 | 93,418,875 (82.6) | 83,383,790 (73.7) | 10,035,085 (8.9) | 34,804,902 (30.8) | 181,597,976 (160.6) |
| Typical-16 | 119,138,942 | 119,073,285 | 101,498,158 (85.2) | 90,320,377 (75.8) | 11,177,781 (9.4) | 40,819,281 (34.3) | 222,431,407 (186.7) |
| Typical-17 | 119,760,200 | 119,661,599 | 100,836,962 (84.2) | 90,348,949 (75.4) | 10,488,013 (8.8) | 36,106,863 (30.1) | 190,751,545 (159.3) |
| Typical-18 | 111,241,066 | 111,139,740 | 96,584,111 (86.8) | 84,083,649 (75.6) | 12,500,462 (11.2) | 44,159,779 (39.7) | 241,863,635 (217.4) |
| Typical-19 | 130,626,166 | 130,517,303 | 110,029,814 (84.2) | 98,256,799 (75.2) | 11,773,015 (9.0) | 44,423,365 (34.0) | 252,761,773 (193.5) |
| Typical-20 | 134,250,278 | 134,146,164 | 113,882,842 (84.8) | 101,889,020 (75.9) | 11,993,822 (8.9) | 41,785,216 (31.1) | 222,678,517 (165.9) |
| Limbic-1 | 128,055,138 | 127,935,605 | 109,770,323 (85.7) | 98,593,301 (77.0) | 11,177,022 (8.7) | 41,791,357 (32.6) | 227,169,478 (177.4) |
| Limbic-2 | 131,223,754 | 131,097,454 | 110,770,095 (84.4) | 98,100,780 (74.8) | 12,669,315 (9.7) | 40,918,851 (31.2) | 228,508,418 (174.1) |
| Limbic-3 | 148,506,328 | 148,345,297 | 125,386,916 (84.4) | 111,145,772 (74.8) | 14,241,144 (9.6) | 47,458,664 (32.0) | 250,901,773 (169.0) |
| Limbic-4 | 122,503,752 | 122,408,327 | 105,169,514 (85.9) | 93,676,424 (76.5) | 11,493,090 (9.4) | 41,497,611 (33.9) | 224,512,856 (183.3) |
| Limbic-5 | 137,500,040 | 137,380,927 | 116,574,780 (84.8) | 107,220,033 (78.0) | 9,354,747 (6.8) | 36,927,991 (26.9) | 193,093,063 (140.4) |
| Limbic-6 | 113,330,370 | 113,226,143 | 96,216,328 (84.9) | 86,551,935 (76.4) | 9,664,393 (8.5) | 36,998,748 (32.6) | 198,689,386 (175.3) |
| Limbic-7 | 103,100,544 | 103,021,768 | 88,905,382 (86.2) | 78,562,694 (76.2) | 10,342,688 (10.0) | 36,865,089 (35.8) | 200,719,088 (194.7) |
| Limbic-8 | 143,963,148 | 143,805,475 | 121,620,275 (84.5) | 110,174,778 (76.5) | 11,445,497 (8.0) | 44,440,332 (30.9) | 242,905,288 (168.7) |
| Limbic-9 | 121,590,604 | 121,472,061 | 106,849,088 (87.9) | 93,720,771 (77.1) | 13,128,317 (10.8) | 44,069,682 (36.2) | 236,142,675 (194.2) |
| Limbic-10 | 137,338,192 | 137,195,789 | 115,045,891 (83.8) | 103,861,577 (75.6) | 11,184,314 (8.1) | 38,278,525 (27.9) | 202,203,461 (147.2) |

### Extended Data Fig. 17

a

| Dataset | Data Type | Description | SynapseID | DoD |
|---|---|---|---|---|
| Mayo RNAseq TCX | Expression | Consensus processed RNASeq raw counts | syn8690799 | 10/2/2019 |
| Mayo RNAseq TCX | Metadata | Individual human and RNAseq | syn3817650 | n/a |
| Mayo RNAseq TCX | Metadata | Quality Control | syn6126114 | n/a |
| MSBB | Expression | Consensus processed RNASeq raw counts | syn8691099 | 10/2/2019 |
| MSBB | Metadata | Individual human | syn6101474 | 11/22/2019 |
| MSBB | Metadata | Assay RNAseq | syn6100548 | 10/2/2019 |

b

| **Dataset** | **Mayo-TCX[h]** | **MSBB-BM22[i]** | **MSBB-BM36[i]** |
|---|---|---|---|
| **Brain Region sampled** | Temporal cortex | Superior temporal gyrus | Parahippocampal gyrus |
| **Unique sample IDs[a]** | 278 | 264 | 267 |
| **Gene counts missing[b]** | 0 | 5 | 4 |
| **Sex check[c]** | 2 | 0 | 0 |
| **RIN <5[d]** | 0 | 38 | 48 |
| **PCA outlier[e]** | 2 | 0 | 0 |
| **Flagged[f]** | 15 | 33 | 48 |
| **Ethnoracial status[g]** | 0 | 39 | 40 |
| **Retained** | 259 | 149 | 127 |
| **AD** | **80** | **70** | **56** |
| **Control** | **68** | **33** | **30** |
| **Other/unknown[j]** | 111 | 46 | 41 |

c

| | **Controls** | **AD** | **p-value** |
|---|---|---|---|
| **Mayo-TCX** | n=68 | n=80 | |
| Age at death, yrs | 86 (78,89) | 85 (78,89) | 0.93 |
| Females, % | 34/68 (50%) | 49/80 (61%) | 0.17 |
| *APOE* ε4, % | 8/68 (12%) | 42/80 (52%) | <0.001 |
| **MSBB-BM22** | n=33 | n=70 | |
| Age at death, yrs | 84 (79,90) | 86 (80,90) | 0.53 |
| Females, % | 18/33 (54%) | 49/70 (70%) | 0.12 |
| *APOE* ε4, % | 2/19 (10%) | 16/44 (36%) | 0.037 |
| **MSBB-BM36** | n=30 | n=56 | |
| Age at death, yrs | 85 (75,90) | 89 (84,90) | 0.16 |
| Females, % | 14/30 (47%) | 41/56 (73%) | 0.015 |
| *APOE* ε4, % | 2/20 (10%) | 9/33 (27%) | 0.13 |

### Extended Data Fig. 18

a

| Antibody | Company | Catalog # | Dilution | Antigen Retrieval | Macro |
|---|---|---|---|---|---|
| **CP13** | Peter Davies gift | n/a | 1:1000 | 30 min. steam in $dH_20$ | CD |
| **Ab39** | Shu-Hui Yen gift | n/a | 1:350 | 30 min. steam in $dH_20$ | PPC |
| **GFAP** | Biogenex | MU020-UC | 1:5000 | 30 min. steam in $dH_20$ | PPC |
| **CD34** | Abcam | Ab81289 | 1:25 | 30 min. steam in $dH_20$ | PPC |
| **CD68** | Dako | M0814 | 1:1000 | 30 min. steam in $dH_20$ | CD |
| **33.1.1** | Pritam Das gift | n/a | 1:1000 | 30 min. in 98% Formic acid followed by 30 min. steam in $dH_20$ | PPC |
| **SERPINA5** | R&D | MAB1266 | 1:100 | 30 min. steam in $dH_20$ | CD |

b

| Primary Antibody | Manufacturer | Concentration | Antigen Retrieval |
|---|---|---|---|
| GFAP | ABCAM ab33922 | 1:800 | 30 min. steam in $dH_20$ |
| MAP2 | ABCAM ab32454 | 1:200 | 30 min. steam in $dH_20$ |
| IBA1 | ABCAM ab178847 | 1:100 | 30 min. steam in citrate |
| CD68 | Cell Signaling 76437 | 1:400 | 30 min. steam in citrate |
| CD34 | ABCAM ab81289 | 1:200 | 30 min. steam in $dH_20$ |
| OLIG2 | ABCAM ab109186 | 1:100 | 30 min. steam in citrate |
| SERPINA5 | R&D MAB1266 | 1:100 | 30 min. steam in $dH_20$ or citrate (refer to co-stain) |
| Tau E1 | Petrucelli Laboratory[1] | 1:1000 | 30 min. steam in $dH_20$ |
| Tau pS396 | Abcam 109390 | 1:1000 | 30 min. steam in $dH_20$ |
| Tau E178 | Abcam 32057 | 1:1000 | 30 min. steam in citrate |
| IgG2a | R&D MAB003 | 1:100 | N/A |

[1]Mayo Clinic, Jacksonville, FL

c

| Secondary Antibody or Stain | Manufacturer | Concentration |
|---|---|---|
| AlexaFluor488 Goat α Mouse | Invitrogen A11001 | 1:500 |
| AlexaFluor568 Goat α Rabbit | Invitrogen A11011 | 1:500 |
| Thioflavin-S | Sigma T1892 | 1% solution dissolved in water |

Extended Data Fig. 19

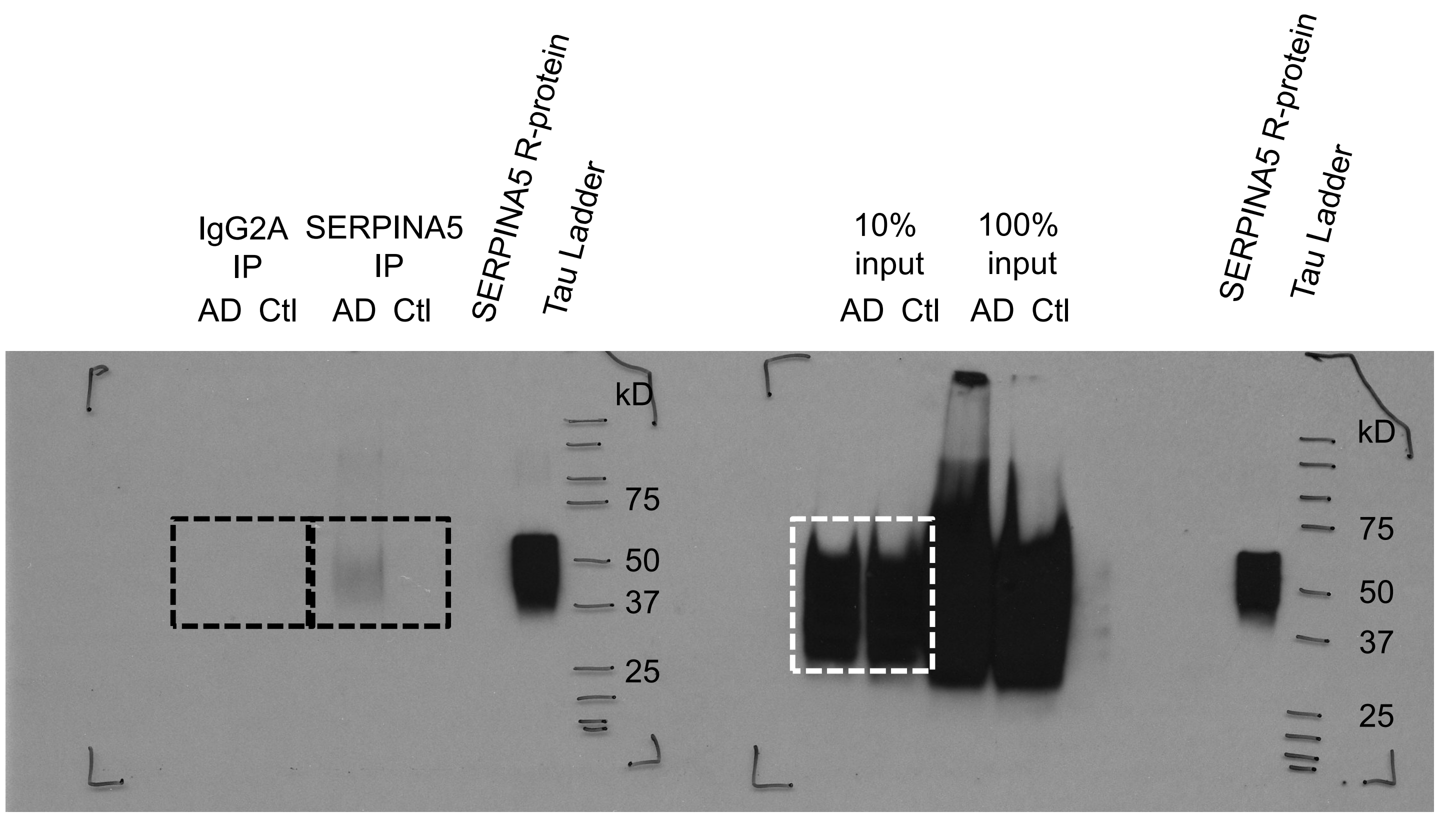

### Extended Data Fig. 20

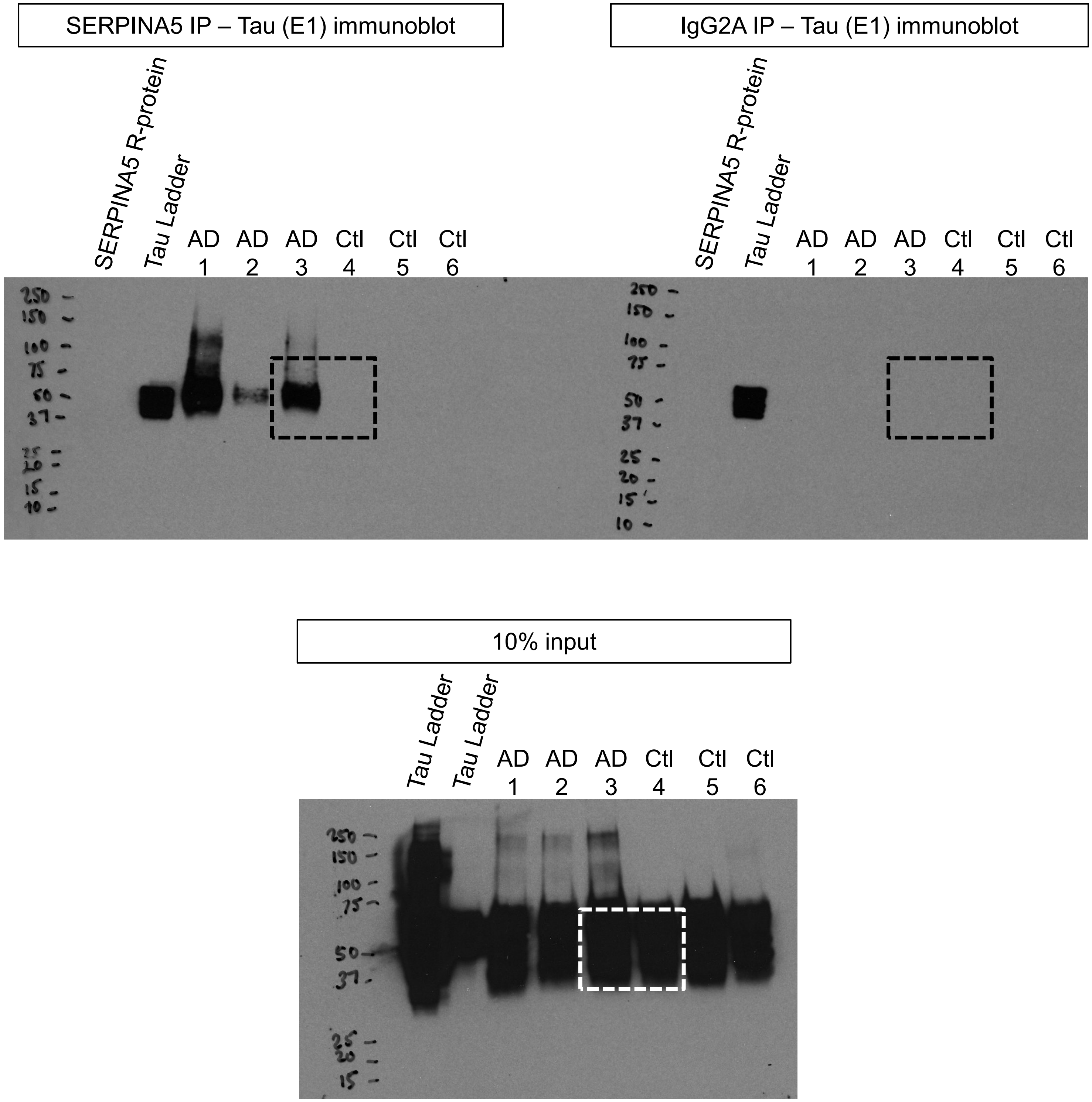
