## Extended Results 1 for "Leveraging selective hippocampal vulnerability among Alzheimer’s disease subtypes reveals a novel tau binding partner SERPINA5"

### Extended Results 1 | Random Forest Importance Measures

| Fig. 4a-b: Control versus all AD subtypes | | | | | | | |
|---|---|---|---|---|---|---|---|
| variable | mean_min_depth | no_of_nodes | accuracy_decrease | gini_decrease | no_of_trees | times_a_root | p_value |
| **SERPINA5** | **2.1797** | **309** | **0.0147** | **3.3182** | **256** | **41** | **<0.0001** |
| **RYBP** | **2.5852** | **246** | **0.0083** | **3.1178** | **209** | **44** | **<0.0001** |
| **SLC38A2** | **2.7758** | **236** | **0.0113** | **3.3811** | **196** | **54** | **<0.0001** |
| **FEM1B** | **3.2232** | **214** | **0.0038** | **1.9006** | **187** | **20** | **<0.0001** |
| **PYDC1** | **3.2656** | **262** | **0.0018** | **1.5229** | **206** | **6** | **<0.0001** |
| RELA | 3.2908 | 209 | 0.0057 | 2.4203 | 182 | 46 | 0.0001 |
| LCOR | 3.3862 | 207 | 0.0051 | 2.2486 | 175 | 37 | 0.0001 |
| DAXX | 3.5147 | 185 | 0.0045 | 2.1183 | 160 | 36 | 0.0232 |
| DNAI1 | 3.6677 | 211 | 0.0029 | 1.5603 | 174 | 3 | <0.0001 |
| OR7A5 | 3.6764 | 191 | 0.0078 | 1.4627 | 170 | 15 | 0.0072 |
| DNAAF1 | 3.782 | 173 | 0.0033 | 1.4835 | 153 | 14 | 0.1436 |
| CTCF | 3.8684 | 173 | 0.0058 | 1.7211 | 151 | 24 | 0.1436 |
| IFITM2 | 3.8903 | 211 | 0.0022 | 1.1026 | 174 | 1 | <0.0001 |
| MAPK15 | 4.0471 | 195 | 0.0027 | 1.1039 | 155 | 0 | 0.003 |
| CAV1 | 4.1811 | 177 | -0.0001 | 0.9606 | 150 | 0 | 0.0845 |
| MAX | 4.1876 | 157 | 0.0037 | 1.1818 | 139 | 11 | 0.5811 |
| NCOA1 | 4.2112 | 167 | 0.0046 | 1.3378 | 138 | 10 | 0.2765 |
| DST | 4.2156 | 154 | 0.003 | 1.2361 | 136 | 16 | 0.6726 |
| RP11.81K2.1 | 4.226 | 150 | 0.0002 | 1.0543 | 125 | 17 | 0.7803 |
| FOXO4 | 4.2947 | 148 | 0.0006 | 0.9129 | 132 | 11 | 0.8256 |
| TAC1 | 4.3413 | 151 | 0.0023 | 0.9947 | 133 | 7 | 0.7553 |
| RBBP7 | 4.3748 | 155 | 0.0019 | 0.8123 | 140 | 1 | 0.6429 |
| LRRC48 | 4.4197 | 155 | 0.0023 | 0.9321 | 132 | 3 | 0.6429 |
| CXCL1 | 4.4378 | 153 | 0.0012 | 0.894 | 138 | 4 | 0.7014 |
| ANGPT2 | 4.4806 | 135 | 0.0032 | 1.0137 | 122 | 14 | 0.9784 |
| DYDC2 | 4.4813 | 150 | 0.002 | 0.961 | 136 | 0 | 0.7803 |
| CNOT8 | 4.5519 | 135 | 0.0035 | 0.7853 | 118 | 7 | 0.9784 |
| RBP1 | 4.5599 | 125 | 0.0003 | 0.6418 | 117 | 1 | 0.998 |
| RAD52 | 4.5845 | 122 | 0.0011 | 0.7876 | 112 | 6 | 0.9992 |
| ALOX15B | 4.6165 | 130 | 0.0021 | 0.808 | 109 | 12 | 0.9928 |
| ERBB2IP | 4.6771 | 129 | 0.0018 | 0.77 | 117 | 2 | 0.9943 |
| BCL2 | 4.7088 | 130 | 0.0021 | 0.8518 | 115 | 13 | 0.9928 |
| PSEN2 | 4.7486 | 126 | 0.0003 | 0.6329 | 112 | 1 | 0.9974 |
| ATR | 4.7665 | 133 | 0.0006 | 0.637 | 119 | 0 | 0.9858 |
| PPM1D | 4.7688 | 117 | -0.0001 | 0.7 | 109 | 4 | 0.9998 |
| MAGED1 | 4.7872 | 126 | 0.001 | 0.6242 | 114 | 2 | 0.9974 |
| SIPA1 | 4.7913 | 125 | 0.0021 | 0.6497 | 113 | 1 | 0.998 |
| INSR | 4.8254 | 111 | 0.0017 | 0.5754 | 101 | 3 | 1 |
| EML4 | 4.8337 | 110 | -0.0003 | 0.5623 | 99 | 1 | 1 |
| CDKN2C | 4.8459 | 113 | 0.0008 | 0.7107 | 97 | 3 | 1 |
| SUN2 | 4.9796 | 102 | 0.0003 | 0.5637 | 93 | 4 | 1 |
| SIRT1 | 4.9902 | 110 | 0.0009 | 0.5715 | 98 | 4 | 1 |
| IRS2 | 5.0862 | 98 | 0.0002 | 0.5796 | 88 | 1 | 1 |
| NOS1 | 5.1311 | 89 | 0 | 0.418 | 80 | 0 | 1 |

| Extended Data Fig. 8a: Representative Phenotype (Control versus Typical AD) | | | | | | | |
|---|---|---|---|---|---|---|---|
| variable | mean_min_depth | no_of_nodes | accuracy_decrease | gini_decrease | no_of_trees | times_a_root | p_value |
| **DNAAF1** | **2.1994** | **189** | **0.0093** | **2.2345** | **169** | **28** | **<0.0001** |
| **SERPINA5** | **2.3181** | **194** | **0.0078** | **1.8661** | **172** | **25** | **<0.0001** |
| **PYDC1** | **2.4054** | **221** | **0.0031** | **1.3465** | **185** | **2** | **<0.0001** |
| **RYBP** | **2.5101** | **178** | **0.008** | **2.1671** | **158** | **37** | **<0.0001** |
| **SLC38A2** | **2.5743** | **164** | **0.0121** | **2.4328** | **142** | **48** | **0.0018** |
| DNAI1 | 2.5764 | 188 | 0.0075 | 1.5829 | 162 | 9 | <0.0001 |
| NCOA1 | 2.5926 | 186 | 0.0096 | 1.7209 | 162 | 15 | <0.0001 |
| FEM1B | 2.848 | 166 | 0.0073 | 1.6579 | 144 | 25 | 0.0011 |
| LCOR | 2.8938 | 154 | 0.0074 | 1.7766 | 133 | 33 | 0.0188 |
| CTCF | 2.9857 | 151 | 0.0096 | 1.8161 | 133 | 33 | 0.0339 |
| DYDC2 | 3.0242 | 177 | 0.0057 | 1.3793 | 149 | 3 | <0.0001 |
| RELA | 3.0288 | 152 | 0.0081 | 1.5441 | 136 | 28 | 0.028 |
| MAPK15 | 3.0806 | 161 | 0.0022 | 1.3591 | 141 | 14 | 0.0039 |
| DAXX | 3.3025 | 124 | 0.0074 | 1.6102 | 113 | 31 | 0.7024 |
| OR7A5 | 3.4147 | 146 | 0.0069 | 1.0304 | 131 | 12 | 0.0807 |
| CAV1 | 3.5064 | 153 | 0.0004 | 0.8063 | 134 | 1 | 0.023 |
| RP11.81K2.1 | 3.5332 | 127 | 0.0058 | 1.1257 | 112 | 22 | 0.6032 |
| TAC1 | 3.6171 | 134 | 0.0024 | 0.945 | 123 | 8 | 0.3595 |
| PSEN2 | 3.6283 | 135 | 0.0026 | 1.0294 | 117 | 13 | 0.3272 |
| ALOX15B | 3.8125 | 123 | 0.0021 | 0.8321 | 111 | 13 | 0.7329 |
| MAGED1 | 3.843 | 124 | 0.0034 | 0.6378 | 113 | 4 | 0.7024 |
| CXCL1 | 3.845 | 120 | 0.0005 | 0.7816 | 111 | 7 | 0.8144 |
| SIPA1 | 3.9122 | 121 | 0.0024 | 0.6509 | 103 | 3 | 0.789 |
| PPM1D | 3.9152 | 123 | 0.0032 | 0.7126 | 111 | 7 | 0.7329 |
| LRRC48 | 3.9464 | 118 | 0.0022 | 0.8817 | 104 | 9 | 0.8593 |
| IFITM2 | 3.9751 | 120 | 0.0011 | 0.5677 | 105 | 0 | 0.8144 |
| EML4 | 4.0078 | 114 | 0.0002 | 0.6264 | 102 | 4 | 0.9259 |
| RBP1 | 4.0205 | 112 | -0.0004 | 0.6672 | 100 | 4 | 0.9486 |
| CNOT8 | 4.0242 | 105 | 0.0013 | 0.6609 | 99 | 7 | 0.989 |
| ANGPT2 | 4.0928 | 107 | 0.0024 | 0.7651 | 98 | 9 | 0.9822 |
| RBBP7 | 4.1972 | 107 | 0.0006 | 0.5529 | 99 | 1 | 0.9822 |
| CDKN2C | 4.1981 | 95 | 0.002 | 0.7694 | 87 | 9 | 0.9994 |
| ERBB2IP | 4.2352 | 103 | 0.0028 | 0.6157 | 96 | 6 | 0.9935 |
| FOXO4 | 4.2506 | 88 | 0.0005 | 0.6509 | 83 | 7 | 1 |
| RAD52 | 4.2573 | 96 | -0.0005 | 0.592 | 90 | 3 | 0.9992 |
| BCL2 | 4.2696 | 103 | 0.0013 | 0.524 | 94 | 2 | 0.9935 |
| MAX | 4.272 | 98 | 0.0019 | 0.6626 | 86 | 5 | 0.9985 |
| INSR | 4.2837 | 108 | 0.0015 | 0.5619 | 99 | 4 | 0.9776 |
| DST | 4.3302 | 88 | 0.0022 | 0.6232 | 79 | 5 | 1 |
| ATR | 4.3581 | 108 | 0.0011 | 0.4717 | 92 | 0 | 0.9776 |
| IRS2 | 4.4921 | 85 | 0.0011 | 0.4139 | 82 | 0 | 1 |
| SUN2 | 4.5425 | 89 | 0.0008 | 0.427 | 83 | 0 | 0.9999 |
| SIRT1 | 4.5613 | 81 | 0.0006 | 0.5007 | 72 | 4 | 1 |
| NOS1 | 4.771 | 66 | -0.0008 | 0.363 | 62 | 0 | 1 |

Extended Data Fig. 8a: Extreme Phenotype (Hippocampal sparing AD versus Limbic predominant AD)

| variable | mean_min_depth | no_of_nodes | accuracy_decrease | gini_decrease | no_of_trees | times_a_root | p_value |
|---|---|---|---|---|---|---|---|
| **FEM1B** | **1.5758** | **228** | **0.03** | **2.9679** | **198** | **59** | **<0.0001** |
| **ERBB2IP** | **2.9319** | **167** | **0.0123** | **1.3556** | **147** | **29** | **<0.0001** |
| **SERPINA5** | **2.9669** | **169** | **0.0045** | **1.404** | **152** | **23** | **<0.0001** |
| **LRRC48** | **3.0664** | **168** | **0.0064** | **1.3735** | **145** | **28** | **<0.0001** |
| **RP11.81K2.1** | **3.294** | **156** | **0.0047** | **1.2357** | **140** | **12** | **0.0004** |
| CAV1 | 3.4272 | 155 | 0.0054 | 1.0236 | 142 | 16 | 0.0006 |
| MAPK15 | 3.4544 | 144 | 0.0047 | 1.2094 | 128 | 28 | 0.0109 |
| RYBP | 3.4791 | 137 | 0.0094 | 1.1538 | 122 | 26 | 0.0466 |
| RAD52 | 3.4945 | 150 | 0.0046 | 1.1268 | 133 | 20 | 0.0024 |
| ATR | 3.5521 | 136 | 0.0005 | 1.057 | 119 | 28 | 0.056 |
| RBP1 | 3.6641 | 134 | 0.0009 | 1.0091 | 120 | 24 | 0.0791 |
| LCOR | 3.774 | 120 | 0.0053 | 0.9222 | 108 | 23 | 0.4454 |
| SIPA1 | 3.8198 | 113 | 0.0022 | 0.9982 | 103 | 32 | 0.6978 |
| CNOT8 | 3.8233 | 128 | 0.0017 | 0.818 | 112 | 15 | 0.1919 |
| BCL2 | 3.846 | 114 | 0.003 | 0.9601 | 104 | 22 | 0.6641 |
| DST | 3.9445 | 123 | 0.0003 | 0.7234 | 112 | 8 | 0.3394 |
| IRS2 | 4.0088 | 131 | -0.0024 | 0.7275 | 116 | 2 | 0.1267 |
| OR7A5 | 4.0451 | 126 | -0.0011 | 0.7052 | 117 | 2 | 0.2454 |
| IFITM2 | 4.0865 | 134 | 0.0028 | 0.769 | 118 | 5 | 0.0791 |
| DAXX | 4.0894 | 111 | 0.0017 | 0.6542 | 105 | 0 | 0.7604 |
| ALOX15B | 4.0957 | 111 | 0.0007 | 0.6706 | 101 | 8 | 0.7604 |
| SLC38A2 | 4.2233 | 108 | -0.0014 | 0.6634 | 97 | 7 | 0.8399 |
| CDKN2C | 4.2473 | 115 | 0.0019 | 0.6659 | 101 | 7 | 0.6291 |
| CTCF | 4.2684 | 113 | 0.0004 | 0.6696 | 102 | 0 | 0.6978 |
| PYDC1 | 4.292 | 122 | 0.0013 | 0.6095 | 111 | 6 | 0.3737 |
| DNAAF1 | 4.3002 | 115 | -0.0007 | 0.5362 | 109 | 2 | 0.6291 |
| PPM1D | 4.3432 | 108 | 0.001 | 0.5471 | 101 | 4 | 0.8399 |
| CXCL1 | 4.361 | 98 | -0.001 | 0.5225 | 93 | 4 | 0.9755 |
| NOS1 | 4.3761 | 105 | -0.0006 | 0.5852 | 93 | 6 | 0.9003 |
| EML4 | 4.3862 | 96 | 0.001 | 0.4927 | 93 | 3 | 0.9849 |
| RELA | 4.4074 | 104 | 0.0013 | 0.5437 | 94 | 0 | 0.9162 |
| FOXO4 | 4.4163 | 91 | -0.0013 | 0.5003 | 82 | 7 | 0.9962 |
| NCOA1 | 4.4491 | 101 | -0.0021 | 0.5644 | 90 | 3 | 0.9528 |
| TAC1 | 4.4825 | 93 | -0.0025 | 0.5216 | 88 | 3 | 0.9932 |
| INSR | 4.5312 | 96 | -0.0007 | 0.5633 | 86 | 5 | 0.9849 |
| SUN2 | 4.5375 | 98 | -0.0006 | 0.5555 | 82 | 7 | 0.9755 |
| MAGED1 | 4.5571 | 87 | 0.0005 | 0.4735 | 76 | 3 | 0.999 |
| MAX | 4.5615 | 88 | 0.0009 | 0.4183 | 86 | 1 | 0.9985 |
| ANGPT2 | 4.5757 | 94 | 0 | 0.4731 | 85 | 2 | 0.991 |
| RBBP7 | 4.6168 | 82 | 0.0011 | 0.4963 | 75 | 4 | 0.9998 |
| SIRT1 | 4.6256 | 87 | -0.0004 | 0.4486 | 79 | 3 | 0.999 |
| PSEN2 | 4.6417 | 89 | -0.0003 | 0.5317 | 80 | 8 | 0.998 |
| DYDC2 | 4.7601 | 81 | -0.0009 | 0.3979 | 77 | 0 | 0.9999 |
| DNAI1 | 4.8302 | 74 | -0.0002 | 0.3635 | 71 | 5 | 1 |

### Column descriptions

| Column | Description |
|---|---|
| accuracy_decrease (classification) | mean decrease of prediction accuracy after X j Xj is permuted |
| gini_decrease (classification) | mean decrease in the Gini index of node impurity (i.e. increase of node purity) by splits on X j Xj |
| mse_increase (regression) | mean increase of mean squared error after X j Xj is permuted |
| node_purity_increase (regression) | mean node purity increase by splits on X j Xj , as measured by the decrease in sum of squares |
| e. mean_minimal_depth | mean minimal depth calculated in one of three ways specified by the parameter mean_sample |
| no_of_trees | total number of trees in which a split on X j Xj occurs |
| no_of_nodes | total number of nodes that use X j Xj for splitting (it is usually equal to no_of_trees if trees are shallow) |
| times_a_root | total number of trees in which X j Xj is used for splitting the root node (i.e., the whole sample is divided into two based on the value of X j Xj ) |
| p_value | p-value for the one-sided binomial test using the following distribution |

For description of other metrics included in these tables, please refer to:
https://cran.r-project.org/web/packages/randomForestExplainer/vignettes/randomForestExplainer.html
