## Extended Results 2 for "Leveraging selective hippocampal vulnerability among Alzheimer’s disease subtypes reveals a novel tau binding partner SERPINA5"

### Extended Results 2 | AMP-AD Top 5 Gene Comparison

**Mayo-TCX: Mayo Clinic Temporal Cortex (syn3163039, TCX)**

| GeneName | log2FC | FDR | p-value | meanCQN.control | meanCQN.AD | AvgMappedReads | Direction |
|---|---|---|---|---|---|---|---|
| SERPINA5 | 2.1 | 0.000040 | 0.00000085 | -0.15 | 1.5 | 103 | Upregulated |
| RYBP | 0.0049 | 1.0 | 0.95 | 5.6 | 5.7 | 3165 | Upregulated |
| SLC38A2 | 1.0 | 0.0000015 | 0.0000000086 | 6.9 | 7.7 | 9841 | Upregulated |
| FEM1B | -0.10 | 0.24 | 0.072 | 6.3 | 6.3 | 5092 | Downregulated |
| PYDC1 | -0.78 | 0.00077 | 0.000039 | 3.2 | 3.2 | 77 | Downregulated |

**MSBB-BM22: Mt. Sinai Brain Bank Superior Temporal Cortex (syn20801188, BM22)**

| GeneName | log2FC | FDR | p-value | meanCQN.control | meanCQN.AD | AvgMappedReads | Direction |
|---|---|---|---|---|---|---|---|
| SERPINA5 | 1.0 | 0.095 | 0.00040 | -0.57 | 0.23 | 21 | Upregulated |
| RYBP | 0.14 | 0.34 | 0.029 | 4.3 | 4.4 | 1024 | Upregulated |
| SLC38A2 | 0.35 | 0.27 | 0.015 | 6.3 | 6.6 | 4105 | Upregulated |
| FEM1B | -0.038 | 0.82 | 0.40 | 6.0 | 6.0 | 3173 | Downregulated |
| PYDC1 | -0.27 | 0.59 | 0.15 | 1.2 | 0.89 | 17 | Downregulated |

**MSBB-BM36: Mt. Sinai Brain Bank Superior Temporal Cortex (syn20801188, BM36)**

| GeneName | log2FC | FDR | p-value | meanCQN.control | meanCQN.AD | AvgMappedReads | Direction |
|---|---|---|---|---|---|---|---|
| SERPINA5 | 2.0 | 0.0016 | 0.0000030 | -1.1 | 0.81 | 31 | Upregulated |
| RYBP | 0.17 | 0.051 | 0.0015 | 4.5 | 4.6 | 1077 | Upregulated |
| SLC38A2 | 0.29 | 0.28 | 0.039 | 6.3 | 6.6 | 3710 | Upregulated |
| FEM1B | -0.10 | 0.21 | 0.022 | 6.2 | 6.0 | 3180 | Downregulated |
| PYDC1 | -0.46 | 0.066 | 0.0024 | 2.2 | 1.7 | 22 | Downregulated |

**Crist et al. (Mayo Clinic Hippocampus)**

| GeneName | log2FC | FDR | p-value | Mean.Ctrl | Mean.Typical | AvgMappedReads | Direction |
|---|---|---|---|---|---|---|---|
| SERPINA5 | 1.7 | 0.0061 | 0.0000041 | -0.87 | 0.68 | 220 | Upregulated |
| RYBP | 0.28 | 0.12 | 0.0027 | 3.2 | 3.4 | 3659 | Upregulated |
| SLC38A2 | 0.93 | 0.0050 | 0.0000026 | 4.7 | 5.5 | 13845 | Upregulated |
| FEM1B | 0.20 | 0.17 | 0.0062 | 4.5 | 4.7 | 9333 | Upregulated |
| PYDC1 | -1.0 | 0.21 | 0.0089 | 0.36 | -0.87 | 18 | Downregulated |
