## Extended Results 3 for "Leveraging selective hippocampal vulnerability among Alzheimer’s disease subtypes reveals a novel tau binding partner SERPINA5"

### Extended Results 3 | Neuner Review[78] Literature Genes

| GeneName | GeneID | Step 4: Group-wise difference controls and AD subtypes p-value must be <0.05 | Step 4: Mono-tonically directed | Step 4: Must be protein coding | Step 4: Neuropathologic measures fdr<0.25, p<0.05 Robustly associate in both analyses | Pass/Fail summary |
|---|---|---|---|---|---|---|
| ABCA7 | ENSG00000064687 | 0.224432 | No | Pass | no association | Fail, group-wise differences p>0.05 |
| ABI3 | ENSG00000108798 | 0.267804 | No | Pass | no association | Fail, group-wise differences p>0.05 |
| AC099552.4 | ENSG00000217825 | 0.484519 | Yes | Fail in Step 0 | no association | Fail in Step 0 (low expressing) |
| ACE | ENSG00000159640 | 0.11499 | Yes | Fail in Step 0 | no association | Fail in Step 0 (low expressing) |
| ACE | ENSG00000264813 | 0.462424 | No | Fail in Step 0 | no association | Fail in Step 0 (low expressing) |
| ACSM1 | ENSG00000166743 | 0.345316 | No | Fail in Step 0 | no association | Fail in Step 0 (low expressing) |
| ADAM10 | ENSG00000137845 | 0.00676065 | Yes | Pass | Ab39 | Passed Step 4 criteria |
| ADAMTS1 | ENSG00000154734 | 0.0281646 | No | Pass | Ab39 | Fail, not monotonically-directed |
| ADAMTS4 | ENSG00000158859 | 0.302511 | No | Pass | no association | Fail, group-wise differences p>0.05 |
| AKAP9 | ENSG00000127914 | 0.322729 | No | Pass | no association | Fail, group-wise differences p>0.05 |
| ALPK2 | ENSG00000198796 | 0.199508 | No | Fail in Step 0 | no association | Fail in Step 0 (low expressing) |
| ANKMY2 | ENSG00000106524 | 0.00086514 | Yes | Pass | Braak; Thal; Ab39 | Passed Step 4 criteria |
| ANKS4B | ENSG00000175311 | 0.0245938 | Yes | Fail in Step 0 | no association | Fail in Step 0 (low expressing) |
| APH1B | ENSG00000138613 | 0.315639 | No | Pass | no association | Fail, group-wise differences p>0.05 |
| APOE | ENSG00000130203 | 0.165588 | No | Pass | no association | Fail, group-wise differences p>0.05 |
| APP | ENSG00000142192 | 0.510598 | No | Pass | no association | Fail, group-wise differences p>0.05 |
| ATP5F1 | ENSG00000116459 | 0.00322088 | Yes | Pass | Braak; Thal; Ab39 | Passed Step 4 criteria |
| B3GALT2 | ENSG00000162630 | 0.345955 | Yes | Pass | no association | Fail, group-wise differences p>0.05 |
| BCKDK | ENSG00000103507 | 0.620264 | Yes | Pass | no association | Fail, group-wise differences p>0.05 |
| BDNF | ENSG00000176697 | 0.234211 | No | Pass | no association | Fail, group-wise differences p>0.05 |
| BIN1 | ENSG00000136717 | 0.320193 | No | Pass | no association | Fail, group-wise differences p>0.05 |
| BLOC1S3 | ENSG00000189114 | 0.254192 | No | Pass | no association | Fail, group-wise differences p>0.05 |
| BZRAP1-AS1 | ENSG00000265148 | 0.415163 | No | Fail | no association | Fail, group-wise differences p>0.05 |
| BZW2 | ENSG00000136261 | 0.154298 | Yes | Pass | no association | Fail, group-wise differences p>0.05 |
| C16orf62 | ENSG00000103544 | 0.51216 | No | Pass | no association | Fail, group-wise differences p>0.05 |
| CASS4 | ENSG00000087589 | 0.0271143 | No | Pass | no association | Fail, not monotonically-directed |
| CD2AP | ENSG00000198087 | 0.817111 | Yes | Pass | no association | Fail, group-wise differences p>0.05 |
| CD33 | ENSG00000105383 | 0.271216 | No | Pass | no association | Fail, group-wise differences p>0.05 |
| CELF1 | ENSG00000149187 | 0.795317 | No | Pass | no association | Fail, group-wise differences p>0.05 |
| CLNK | ENSG00000109684 | 0.310674 | No | Fail in Step 0 | no association | Fail in Step 0 (low expressing) |
| CLU | ENSG00000120885 | 0.215988 | Yes | Pass | no association | Fail, group-wise differences p>0.05 |
| CNTN5 | ENSG00000149972 | 0.120932 | No | Pass | no association | Fail, group-wise differences p>0.05 |
| CNTNAP2 | ENSG00000174469 | 0.19226 | Yes | Pass | no association | Fail, group-wise differences p>0.05 |
| COBL | ENSG00000106078 | 0.770243 | No | Pass | no association | Fail, group-wise differences p>0.05 |
| CR1 | ENSG00000203710 | 0.397434 | No | Fail in Step 0 | no association | Fail in Step 0 (low expressing) |
| CTDP1 | ENSG00000060069 | 0.102563 | No | Pass | no association | Fail, group-wise differences p>0.05 |
| DAPK1 | ENSG00000196730 | 0.140857 | No | Pass | no association | Fail, group-wise differences p>0.05 |
| DSG2 | ENSG00000046604 | 0.249527 | Yes | Fail in Step 0 | no association | Fail in Step 0 (low expressing) |
| ECHDC3 | ENSG00000134463 | 0.393534 | No | Pass | no association | Fail, group-wise differences p>0.05 |
| EPHA1 | ENSG00000146904 | 0.131115 | Yes | Fail in Step 0 | no association | Fail in Step 0 (low expressing) |
| EXOC3L2 | ENSG00000130201 | 0.617172 | No | Fail in Step 0 | no association | Fail in Step 0 (low expressing) |
| FAM63B | ENSG00000128923 | 0.292611 | No | Pass | no association | Fail, group-wise differences p>0.05 |
| FBP1 | ENSG00000165140 | 0.80813 | No | Pass | no association | Fail, group-wise differences p>0.05 |
| FERMT2 | ENSG00000073712 | 0.644659 | No | Pass | no association | Fail, group-wise differences p>0.05 |
| FRA10AC1 | ENSG00000148690 | 0.269099 | No | Pass | no association | Fail, group-wise differences p>0.05 |

| GeneName | GeneID | Step 4: Group-wise difference controls and AD subtypes p-value must be <0.05 | Step 4: Mono-tonically directed | Step 4: Must be protein coding | Step 4: Neuropathologic measures fdr<0.25, p<0.05 Robustly associate in both analyses | Pass/Fail summary |
|---|---|---|---|---|---|---|
| GALNT7 | ENSG00000109586 | 0.888093 | No | Pass | no association | Fail, group-wise differences p>0.05 |
| GDE1 | ENSG00000006007 | 0.302725 | No | Pass | no association | Fail, group-wise differences p>0.05 |
| GLIS1 | ENSG00000174332 | 0.143503 | Yes | Pass | no association | Fail, group-wise differences p>0.05 |
| GLIS3 | ENSG00000107249 | 0.237899 | No | Pass | no association | Fail, group-wise differences p>0.05 |
| GMNC | ENSG00000205835 | 0.297753 | Yes | Pass | no association | Fail, group-wise differences p>0.05 |
| GPRC5B | ENSG00000167191 | 0.0518086 | Yes | Pass | Ab39 | Fail, group-wise differences p>0.05 |
| GRIN2B | ENSG00000273079 | 0.398606 | No | Pass | no association | Fail, group-wise differences p>0.05 |
| HBEGF | ENSG00000113070 | 0.982263 | No | Pass | no association | Fail, group-wise differences p>0.05 |
| HESX1 | ENSG00000163666 | 0.57992 | No | Pass | no association | Fail, group-wise differences p>0.05 |
| HLA-DRB1 | ENSG00000196126 | 0.114551 | No | Pass | no association | Fail, group-wise differences p>0.05 |
| HLA-DRB5 | ENSG00000198502 | 0.863747 | No | Pass | no association | Fail, group-wise differences p>0.05 |
| HS3ST1 | ENSG00000002587 | 0.0416514 | No | Pass | no association | Fail, not monotonically-directed |
| IL34 | ENSG00000157368 | 0.419555 | No | Pass | no association | Fail, group-wise differences p>0.05 |
| INPP5D | ENSG00000168918 | 0.667486 | No | Pass | no association | Fail, group-wise differences p>0.05 |
| IQCK | ENSG00000174628 | 0.596658 | Yes | Pass | no association | Fail, group-wise differences p>0.05 |
| ITGA8 | ENSG00000077943 | 0.00867443 | No | Pass | Thal | Fail, not monotonically-directed |
| KAT8 | ENSG00000103510 | 0.0231268 | Yes | Pass | no association | Fail, no association with neuropath |
| KCNH6 | ENSG00000173826 | 0.395575 | No | Fail in Step 0 | no association | Fail in Step 0 (low expressing) |
| KNOP1 | ENSG00000103550 | 0.125756 | No | Pass | no association | Fail, group-wise differences p>0.05 |
| LRP6 | ENSG00000070018 | 0.659092 | No | Pass | no association | Fail, group-wise differences p>0.05 |
| MARK4 | ENSG00000007047 | 0.992413 | No | Pass | no association | Fail, group-wise differences p>0.05 |
| MEF2C | ENSG00000081189 | 0.158461 | No | Pass | no association | Fail, group-wise differences p>0.05 |
| MS4A2 | ENSG00000149534 | 0.868523 | No | Fail in Step 0 | no association | Fail in Step 0 (low expressing) |
| MS4A4A | ENSG00000110079 | 0.802155 | No | Pass | no association | Fail, group-wise differences p>0.05 |
| MS4A6A | ENSG00000110077 | 0.097507 | No | Pass | no association | Fail, group-wise differences p>0.05 |
| MS4A6E | ENSG00000166926 | 0.0317935 | No | Fail in Step 0 | no association | Fail in Step 0 (low expressing) |
| NFATC1 | ENSG00000131196 | 0.919537 | Yes | Pass | no association | Fail, group-wise differences p>0.05 |
| NME8 | ENSG00000086288 | 0.23112 | No | Fail in Step 0 | no association | Fail in Step 0 (low expressing) |
| NOTCH3 | ENSG00000074181 | 0.755962 | No | Pass | no association | Fail, group-wise differences p>0.05 |
| NYAP1 | ENSG00000166924 | 0.963773 | No | Pass | no association | Fail, group-wise differences p>0.05 |
| OARD1 | ENSG00000124596 | 0.286194 | No | Pass | no association | Fail, group-wise differences p>0.05 |
| OSTN | ENSG00000188729 | 0.168762 | No | Fail in Step 0 | no association | Fail in Step 0 (low expressing) |
| PCDH8 | ENSG00000136099 | 0.549915 | No | Pass | no association | Fail, group-wise differences p>0.05 |
| PFDN1 | ENSG00000113068 | 0.0902279 | No | Pass | no association | Fail, group-wise differences p>0.05 |
| PICALM | ENSG00000073921 | 0.254338 | No | Pass | no association | Fail, group-wise differences p>0.05 |
| PILRA | ENSG00000085514 | 0.923296 | No | Pass | no association | Fail, group-wise differences p>0.05 |
| PIP | ENSG00000159763 | 0.559166 | No | Fail in Step 0 | no association | Fail in Step 0 (low expressing) |
| PLCG2 | ENSG00000197943 | 0.370988 | No | Pass | no association | Fail, group-wise differences p>0.05 |
| PLD3 | ENSG00000105223 | 0.599286 | No | Pass | no association | Fail, group-wise differences p>0.05 |
| PRNP | ENSG00000171867 | 0.666178 | No | Pass | no association | Fail, group-wise differences p>0.05 |
| PSEN1 | ENSG00000080815 | 0.79361 | No | Pass | no association | Fail, group-wise differences p>0.05 |
| PSEN2 | ENSG00000143801 | 0.0154051 | Yes | Pass | Thal; Ab39 | Passed Step 4 criteria |
| PSMB8 | ENSG00000204264 | 0.200604 | No | Pass | no association | Fail, group-wise differences p>0.05 |
| PTK2B | ENSG00000120899 | 0.487029 | No | Pass | no association | Fail, group-wise differences p>0.05 |
| PVRL2 | ENSG00000130202 | 0.77427 | Yes | Pass | no association | Fail, group-wise differences p>0.05 |
| RBFOX1 | ENSG00000078328 | 0.526243 | No | Pass | no association | Fail, group-wise differences p>0.05 |
| RIN3 | ENSG00000100599 | 0.0346903 | No | Pass | no association | Fail, not monotonically-directed |
| SCIMP | ENSG00000161929 | 0.775518 | Yes | Fail in Step 0 | no association | Fail in Step 0 (low expressing) |

| GeneName | GeneID | Step 4: Group-wise difference controls and AD subtypes p-value must be <0.05 | Step 4: Mono-tonically directed | Step 4: Must be protein coding | Step 4: Neuropathologic measures fdr<0.25, p<0.05 Robustly associate in both analyses | Pass/Fail summary |
|---|---|---|---|---|---|---|
| SERPINB1 | ENSG00000021355 | 0.858301 | No | Pass | no association | Fail, group-wise differences p>0.05 |
| SLC10A2 | ENSG00000125255 | 0.636976 | No | Fail in Step 0 | no association | Fail in Step 0 (low expressing) |
| SLC24A4 | ENSG00000140090 | 0.252707 | Yes | Pass | no association | Fail, group-wise differences p>0.05 |
| SLTM | ENSG00000137776 | 0.280178 | No | Pass | no association | Fail, group-wise differences p>0.05 |
| SORL1 | ENSG00000137642 | 0.0109846 | No | Pass | no association | Fail, not monotonically-directed |
| SPI1 | ENSG00000066336 | 0.145055 | No | Pass | no association | Fail, group-wise differences p>0.05 |
| SPPL2A | ENSG00000138600 | 0.467655 | No | Pass | no association | Fail, group-wise differences p>0.05 |
| STYX | ENSG00000198252 | 0.22876 | No | Pass | no association | Fail, group-wise differences p>0.05 |
| TAP2 | ENSG00000204267 | 0.0177813 | No | Pass | no association | Fail, not monotonically-directed |
| TF | ENSG00000091513 | 0.848483 | No | Pass | no association | Fail, group-wise differences p>0.05 |
| TOMM40 | ENSG00000130204 | 0.11573 | No | Pass | no association | Fail, group-wise differences p>0.05 |
| TREM2 | ENSG00000095970 | 0.620013 | No | Pass | no association | Fail, group-wise differences p>0.05 |
| UNC5C | ENSG00000182168 | 0.628214 | No | Pass | no association | Fail, group-wise differences p>0.05 |
| USP6NL | ENSG00000148429 | 0.0676954 | No | Pass | no association | Fail, group-wise differences p>0.05 |
| WWOX | ENSG00000186153 | 0.632314 | No | Pass | no association | Fail, group-wise differences p>0.05 |
| ZNF232 | ENSG00000167840 | 0.151841 | No | Pass | no association | Fail, group-wise differences p>0.05 |
| SUZ12P1 | -- | -- | -- | -- | -- | Not identified in our dataset |
| PPB | -- | -- | -- | -- | -- | Not identified in our dataset |
| NCR2 | -- | -- | -- | -- | -- | Not identified in our dataset |
| BZRAP-AS1 | -- | -- | -- | -- | -- | Not identified in our dataset |
